# Quantitative comparison of principal component analysis and unsupervised deep learning using variational autoencoders for shape analysis of motile cells

**DOI:** 10.1101/2020.06.26.174474

**Authors:** Caleb K. Chan, Amalia Hadjitheodorou, Tony Y.-C. Tsai, Julie A. Theriot

**Author notes:** <u>Corresponding Author:</u> Julie A. Theriot, Ph.D., Department of Biology, University of Washington, Life Sciences Building, Room 575, Box 351800, Seattle, WA 98195-1800, USA.

## Abstract

Cell motility is a crucial biological function for many cell types, including the immune cells in our body that act as first responders to foreign agents. In this work we consider the amoeboid motility of human neutrophils, which show complex and continuous morphological changes during locomotion. We imaged live neutrophils migrating on a 2D plane and extracted unbiased shape representations using cell contours and binary masks. We were able to decompose these complex shapes into low-dimensional encodings with both principal component analysis (PCA) and an unsupervised deep learning technique using variational autoencoders (VAE), enhanced with generative adversarial networks (GANs). We found that the neural network architecture, the VAE-GAN, was able to encode complex cell shapes into a low-dimensional latent space that encodes the same shape variation information as PCA, but much more efficiently. Contrary to the conventional viewpoint that the latent space is a “black box”, we demonstrated that the information learned and encoded within the latent space is consistent with PCA and is reproducible across independent training runs. Furthermore, by including cell speed into the training of the VAE-GAN, we were able to incorporate cell shape and speed into the same latent space. Our work provides a quantitative framework that connects biological form, through cell shape, to a biological function, cell movement. We believe that our quantitative approach to calculating a compact representation of cell shape using the VAE-GAN provides an important avenue that will support further mechanistic dissection of cell motility.

**AUTHOR SUMMARY:** Deep convolutional neural networks have recently enjoyed a surge in popularity, and have found useful applications in many fields, including biology. Supervised deep learning, which involves the training of neural networks using existing labeled data, has been especially popular in solving image classification problems. However, biological data is often highly complex and continuous in nature, where prior labeling is impractical, if not impossible. Unsupervised deep learning promises to discover trends in the data by reducing its complexity while retaining the most relevant information. At present, challenges in the extraction of meaningful human-interpretable information from the neural network’s nonlinear discovery process have earned it a reputation of being a “black box” that can perform impressively well at prediction but cannot be used to shed any meaningful insight on underlying mechanisms of variation in biological data sets. Our goal in this paper is to establish unsupervised deep learning as a practical tool to gain scientific insight into biological data by first establishing the interpretability of our particular data set (images of the shapes of motile neutrophils) using more traditional techniques. Using the insight gained from this as a guide allows us to shine light into the “black box” of unsupervised deep learning.

## INTRODUCTION

Morphometrics, the measurement of shape, is a fundamental method in biology, with widespread application in studies of taxonomy, development and evolution (1–3), because of the deep connections between form and function in living organisms (4). In cell biology, it is possible to recognize most functionally distinct cell types and determine their physiological state by direct examination of their microscopic organization alone (5,6). However, it is challenging to represent variations in cell shape and organization within a quantitative, objective framework that is directly interpretable by cell biologists. From microscopy images, it is possible to calculate hundreds of objectively measureable image features, including cell properties such as size, aspect ratio and asymmetry, as well as many specific features measuring distributions and textures of fluorescent protein labels (7,8). These kinds of “high-content” approaches to image analysis have proved to be useful for applications such as drug screening and phenotype evaluation for large-scale genetic perturbation experiments, but the highly multidimensional nature of these measurements effectively precludes their direct utility for developing insight into the underlying nature of quantitative variation in cellular morphology. To this end, multiple approaches have been developed to reduce the dimensionality of cell image data to retain meaningful information in a compact space that may be interpreted by inspection (9,10).

For example, principal component analysis (PCA) is an unsupervised, data-driven linear transformation that identifies orthogonal dimensions in a data set containing the maximum amount of variation. Intuitively, the first axis, or principal mode, spans the direction along which the data is most spread out, or contains the most information. The second principal mode spans the direction (orthogonal to the first) with the second largest amount of variation, and so on. By retaining only the top-ranked principal modes that contain the most information, a compact representation of cell shape can therefore be created. Depending on the number of principal modes retained, cell shapes can be reconstructed with any chosen level of accuracy (9).

Because of its simple and intuitively understandable properties, PCA has proved to be valuable for simplifying representations of complex cell image data in a way that can facilitate generation of testable hypotheses relevant to understanding the underlying molecular mechanisms determining cell shape. For example, crawling or amoeboid motility of eukaryotic cells depends on interactions between the dynamic force-generating cytoskeleton and the cell’s plasma membrane (11). Therefore, direct examination of the principal modes of overall cell shape variation in motile cells such as fish epidermal keratocytes and the cellular slime mold *Dictyostelium discoideum* have revealed useful insights into mechanisms governing cell movement speed, balance between adhesion and contractility, and chemotactic behavior (12–14). Similarly, principal modes of shape variation in the bacterium *Caulobacter crescentus* can be directly mapped onto protein functions identified by specific point mutations in the shape-determining protein MreB (15).

While linear dimensionality reduction methods such as PCA are widely used in cell biology, nonlinear methods have not been as well explored. In many cases, nonlinear methods may be able to represent biological shapes more accurately than PCA in a smaller number of dimensions, simplifying their interpretation (16). One particularly intriguing nonlinear technique that is also capable of projecting high-dimensional data into a low-dimensional space is the variational autoencoder (VAE) (17). This unsupervised deep learning framework involves the training of a convolutional neural network (CNN), called the encoder, to encode input images into a compact, low dimensional space (the latent space), along with a second CNN, the decoder, that attempts to reconstruct the input image based only on the compact information stored in the latent space. In order to retain the maximum amount of useful information and maintain pre-existing relationships between data points, the autoencoder is trained to reproduce the input images as accurately as possible while using the low dimensionality of the latent space as a throttle. The number of latent space dimensions of the VAE can be arbitrarily defined prior to training, and the lower the number of latent space dimensions, the more compact the encoding. The optimal number of dimensions is based on the concept of Shannon’s entropy (18,19), which specifies the minimal resource required to retain maximal useful information by utilizing the likelihood of each event’s occurrence. This is analogous to the process of determining how many major principal components to retain in PCA. In fact, it has been shown mathematically that when an autoencoder is constructed using only linear components, the latent space dimensions will approximate the principal modes of PCA (20,21). However, if this restriction is lifted, the autoencoder can also perform dimensionality reduction for data sets with non-linear properties (17,22).

In this paper, we used a variation of the VAE, called the VAE-GAN, that employs two generative adversarial networks (GANs) (23). In this architecture, one GAN is used to enforce a prior normal distribution onto the trained latent space, and the second is devoted to improving the reconstruction quality of the decoder to generate “photorealistic” synthetic images that are highly similar to the real input images (24). We wished to determine whether the VAE-GAN could encode cell shape information for motile cells in a way that might be comparable to the interpretable principal shape modes discoverable by PCA.

To this end, we assembled a dataset consisting of time-lapse videomicroscopy sequences of live crawling human neutrophils. As first responders to sites of infection, the neutrophils’ migration abilities are closely coupled to their physiological function of tracking down and internalizing invading microbes (25). Neutrophils are highly dynamic cells that use a variety of strategies to enter and efficiently maneuver through a range of complex physiological environments (26,27). Their rapid migration is characterized by an amoeboid style of locomotion, featuring dynamic pseudopod protrusion and retraction, flexible oscillatory shape changes, and low adhesion affinity to their substrate (26,28). When confined to a quasi-two dimensional environment, they are able to vary their movement speed and direction, and also change shape dramatically, while maintaining a remarkably constant projected area (28). Therefore, these cells represent an ideal test case for examining whether and how changes in cell shape might be correlated with changes in movement behavior. We included in our data set both primary human neutrophils isolated from fresh blood (human polymorphonuclear leukocytes, or PMNs) and differentiated immortalized HL60 cells, originally derived from a patient with acute promyelocytic leukemia (29), which have been widely used as a genetically manipulable model system for neutrophil migration (28, 30, 31) and immune function (32).

We first generated a numerical representation of cell shape using parameterized 2D cell contours, followed by a statistical analysis using this representation with PCA. We then compared the shape space of our PCA to the latent space of a VAE-GAN trained on binary mask images of the same data set, and found that the shape variations extracted by both techniques are comparable, although the VAE-GAN can achieve better reconstruction accuracy with a substantially more compact representation than PCA. Because CNN architectures are notorious for producing very different results when retrained using the same data sets with different random number seeds or input order (33), we also explicitly examined the reproducibility of VAE-GAN latent spaces retrained independently multiple times, and found that their reproducibility was quite good, particularly when the latent space was restricted to a very small number of dimensions. Finally, we integrated our findings on neutrophil shape changes with motility behavior by including both cell shape and speed information in our input data set for the training of VAE-GAN latent spaces. With that, we constructed a predictive framework using linear regression models based on these new latent space encodings, which allowed us to perform bidirectional predictions between cell shape and speed. The ease and flexibility of incorporation of different types of biological information within a single trained latent space highlights a particular advantage of the VAE-GAN. The combination of shape and cell speed illustrates only one such example, and we believe further applications of this framework will facilitate hypothesis generation and dissection of the mechanisms that support cell motility.

## RESULTS

### PCA decomposes neutrophil cell shapes into biologically interpretable shape modes of motility behavior

Both PMNs and HL60s exhibit dramatic shape changes during migration on glass coverslips under an agarose overlay (28). We collected time-lapse movies with various time intervals and various overall durations using phase-contrast microscopy, resulting in a total data set of 1298 cell images for PMNs and 1715 images for HL60 cells. In order to establish a PCA-based cell shape space, we first aligned all movies to be centered at the centroid of the cell nucleus, cropped the images to a consistent size, segmented the images to obtain a binary mask of the cell outline, and rotated them so that the instantaneous direction of migration was oriented toward the right (Fig. S1A; see *Methods*). Representative images of each cell type after alignment and segmentation are shown in Fig. 1A, and representative time-lapse movies in Supplementary Movie S1. Next, we represented each two-dimensional segmented cell shape as a closed, parameterized contour using a set of 300 coordinates [*x(p), y(p)*], such that each cell shape can be represented as a single point in a 600-dimensional space (Fig. S1B). A slight difference between cell shapes of the two cell types can be observed by comparing their average shapes (Fig. 1A, right).

**Figure 1.**
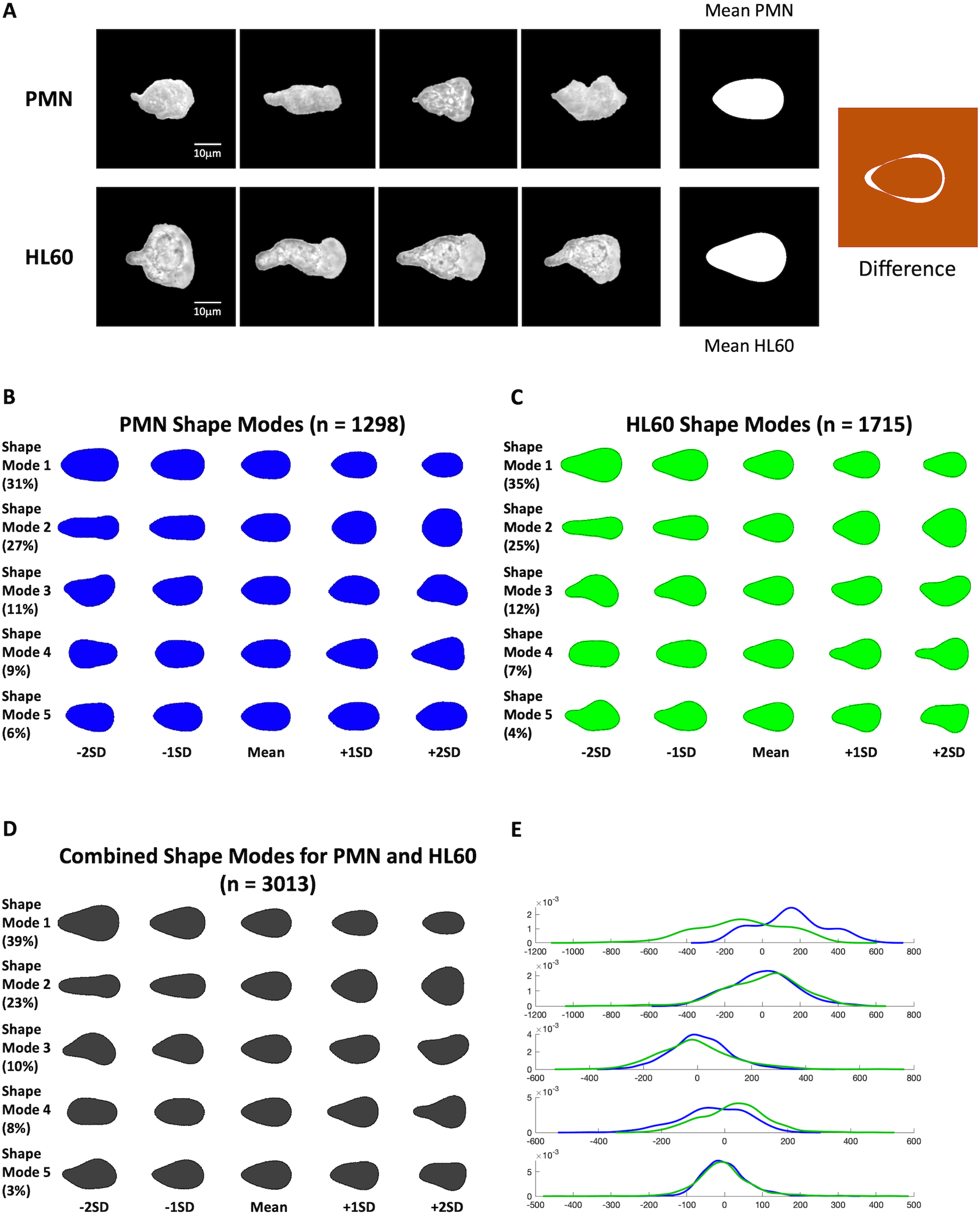
PCA analysis of PMN and HL60 cell shapes. (A) Left, representative images of individual motile cell shapes for PMNs (top) and HL60s (bottom). All images have been centered on the nucleus centroid and rotated so that the direction of movement is toward the right, and cell outlines have been segmented out. Right, average (mean) shapes for PMNs (top) and HL60s (bottom). Far right, difference map for PMN vs. HL60 mean shapes. (B) Top five principal shape modes for PMN-only data set, derived from PCA. (C) Top five principal shape modes for HL60-only data set. (D) Top five principal shape modes for combined PMN and HL60 data set. (E) Probability density estimates of PC scores for PMNs (blue) and HL60s (green) plotted separately for each of the top five principal modes in the combined data set.

By performing PCA on contours for each cell type separately, we observed similar kinds of shape variations over the top five principal shape modes (Fig. 1B-C). Each of the orthogonal shape modes can be readily interpreted as representing a distinct kind of expected biological variation among the cell images. For both cell types, shape mode 1 roughly corresponds to cell size, and shape mode 2 roughly corresponds to cell aspect ratio, analogous to the first two principal shape modes for a different motile cell type, the fish epidermal keratocyte (12). The third principal shape mode represents left-right asymmetry consistent with shapes of cells smoothly turning clockwise or counterclockwise (28), and the fourth represents variations in front-rear asymmetry consistent with shape changes reflecting polarization toward the direction of motion (34). The fifth mode appears to represent a distinct kind of left-right asymmetry. These top five major shape modes explained most of the variations within both data sets (84% for the PMN-only data set and 83% for the HL60-only data set). Thus, focusing on the top five PCA shape modes results in a dramatic reduction of dimensionality (from 600 down to 5), while preserving most of the information regarding cell shape variation. When we combined the two cell types to perform a joint PCA, we saw that the top shape modes and their respective rankings, as well as the motility behaviors apparently described by them, were preserved (Fig. 1D).

Next, we asked whether we can separate the two cell types of the combined data set. In the joint PCA, we observed moderate group separation between the two cell types, as shown by the probability density estimates of the PCA scores for each shape mode, color-coded by cell type (Fig. 1E). The major determinant in PCA’s ability to distinguish between these two cell types was mostly based on shape mode 1 (cell area), although shape mode 4 (degree of polarization) also seemed to contribute to a lesser degree.

We then set out to evaluate the reconstruction accuracy that can be achieved by using information retained in the top PCA shape modes. In principle, a cell’s shape can be reconstructed with 100% accuracy if the PCA scores from all available shape modes are used. Here, for any particular cell within the data set, reconstruction accuracy increased with an increasing number of shape modes used for reconstruction, as expected (Fig. 2A). Fig. 2A shows both the reconstructed images of a single cell as well as the differences between the reconstructed images and the original binary mask representations when the first 2, 3, 4, 8, 16, and 32 shape modes were used. Using reconstruction error as a metric to measure reconstruction accuracy (see *Methods*) and plotting the average reconstruction error for a test set of cell shapes that were held out from the initial set used to calculate the principal shape modes (n = 159), we also saw quantitatively that reconstruction error decreased as we included more shape modes in the reconstruction (Fig. 2B).

**Figure 2.**
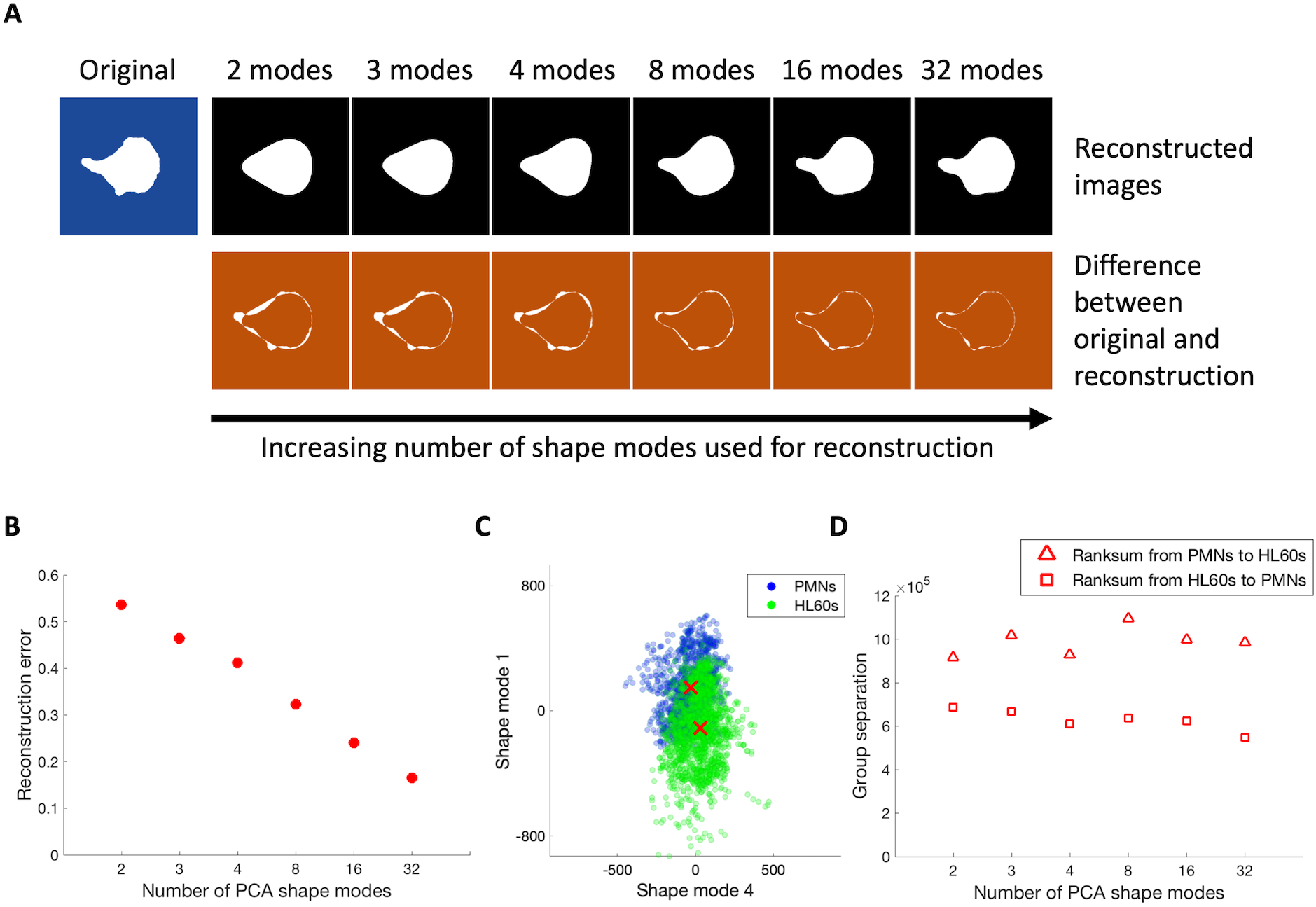
Cell shape reconstruction using different number of PCA shape modes. (A) Cell shape reconstruction of a single cell using 2 to 32 shape modes. Original binary mask is shown in blue, reconstructions in black, and difference maps comparing the reconstructed images to the original binary mask in orange. (B) Average PCA reconstruction error for all cells from 2 to 32 shape modes. (C) Scatter plot showing group separation between PMN and HL60 cells using principal shape modes 1 & 4 (red crosses = median points for each group). (D) Measure of PCA group separation ability using 2 to 32 shape modes.

We noted earlier that the two cell types represented in our data set (PMNs and HL60s) exhibit subtle shape differences, as illustrated by the difference between their average shapes during motility (Fig. 1A). The effectiveness of a shape representation lies not only in the ability to generate the most compact encoding possible, but also in the ability to retain biologically important information that allows for the discrimination between groups carrying distinct phenotypes. Therefore, by including two cell lines that are functionally similar, but with slight phenotypic differences in terms of cell shape, we can quantitatively assess the sensitivity of PCA in terms of its ability to separate the subtle yet observable differences between the two cell types. Fig. 2C shows a sample two-dimensional plot for the PC scores of shape mode 1 versus shape mode 4. By coloring the two cell types differently (PMNs in blue and HL60s in green) we observed separation between the overlapping data points of the two groups, as indicated by their respective medians (red cross symbols) (Fig. 2C). Group separation between the two cell types was quantified using a non-parametric Wilcoxon rank-sum test that measures the degree of overlap between two sets of N-dimensional data points. If two groups of data points were completely separated with no overlap, the metric would return a zero value. On the other hand, the metric would return a large value if two sets of data points were completely overlapping each other. When we calculated group separation using the PC scores from the first 2, 3, 4, 8, 16, and 32 shape modes, we found that the calculated group separation values remained quite stable moving from 2 to 32 shape modes (Fig. 2D).

In summary, by performing PCA on cell shapes for crawling human neutrophils, we found that: 1) the first five major shape modes explained more than 80% of the variations in the data set, 2) the same five shape modes were also biologically interpretable as expected cell-to-cell variations that are experimentally observable, 3) the types and ranking of variations for PMNs as compared to HL60s were the same for the first five major modes, 4) reconstruction error decreased as the number of shape modes used for reconstruction increased, and 5) PCA of cell contours can discriminate between the two cell types.

### VAE-GAN achieves higher reconstruction accuracy than PCA with comparable group separation power

Another approach to reduce the dimensionality of data, analogous to PCA, is the VAE-GAN, which can also encode shape information into compact representations within a low-dimensional latent space (17,24). Therefore, using reconstruction error as a metric again, we asked whether the encoding efficiencies are comparable between these two techniques in terms of their ability to accurately reconstruct the original cell shapes from their compact encodings, while preserving relevant biological information.

The training of a VAE-GAN (Fig. S1C) is a data-driven process. The VAE-GAN consists of two convolutional neural networks that are connected back-to-back, where the goal of the first network, called the encoder, is to take the input images (in our case are the binary masks) and learn to extract the relevant common features of the images. Images with similar features are then embedded into the latent space as points that are close to each other, while images with different features are mapped further apart. The second network then tries to reconstruct the original images based on the latent space encodings learned by the first network, and, through an iterative training process that minimizes the reconstruction error, the two networks are iteratively optimized to allow reconstruction of the data as accurately as possible. The optimization process also includes two additional neural networks (GANs), one to enforce a Gaussian distribution for all dimensions on the encoding of data points in the latent space, and another to ensure that the output images are “photorealistic” and closely resemble the input images (23,24). After the training has converged, a compact representation of cell shapes can be obtained from the latent space where each image is encoded as a low dimensional vector.

The number of dimensions occupied by the latent space is a network architecture-level parameter that is set prior to the training process. Depending on the size of the data set and the complexity of the input image features, the optimal number of latent space dimensions is empirically determined. In general, the number of latent space dimensions sets the degree of dimensionality reduction of the input data and reflects the tradeoff between having a more compact representation of the input data versus having more dimensions to retain additional information about the data set.

Here we considered the same number of latent space dimensions (2, 3, 4, 8, and 16) as the number of shape modes used to evaluate our PCA’s reconstruction accuracy, as a fair comparison. In our PCA methodology, about 80% of the data set’s variations could be explained using the first five major shape modes, and the image reconstruction accuracy steadily improved by incorporating more shape modes. Therefore, we set out to compare the image reconstruction accuracy between the PCA methodology and the VAE-GAN using 2-16 shape modes or latent space dimensions.

For each of the latent space dimensions considered, we trained the VAE-GAN using the same training data set we used for PCA. We then applied the same test data set used previously for PCA, and compared the image reconstruction accuracy between the VAE-GAN and PCA for equivalent numbers of dimensions. For any specific number of dimensions, we found that the reconstructed shapes generated by the VAE-GAN were much more similar to the original image as compared to the PCA output. This was especially noticeable when using a low number of latent space dimensions, for instance, when using only 2 (Fig. 3A). We then compared quantitatively the reconstruction errors between the VAE-GAN and PCA using the same test data set and the same number of dimensions as shape modes from PCA (Fig. 3B). The VAE-GAN reconstruction results were averaged from 4 independent training sessions using different initial training parameters. Consistent with our qualitative visual observation from Fig. 3A, we found that the reconstruction error of the VAE-GAN is at least 5-fold smaller than that of PCA at N = 2, although that gap narrowed as the number of dimensions increased. For PCA the reconstruction error decreased by about 2-fold when going from 2 to 16 shape modes, from over 0.5 down to about 0.25 (Fig. 2B), while the VAE-GAN continued to maintain a reconstruction error of less than 0.1 throughout (Fig. 3B). Impressively, more than 512 PCA shape modes were required to approach a reconstruction error of 0.08, which is about the same as the VAE-GAN reconstruction error at just N = 2. It is clear that the image reconstruction accuracy of the VAE-GAN is vastly superior to that of PCA in low-dimensional shape space.

**Figure 3.**
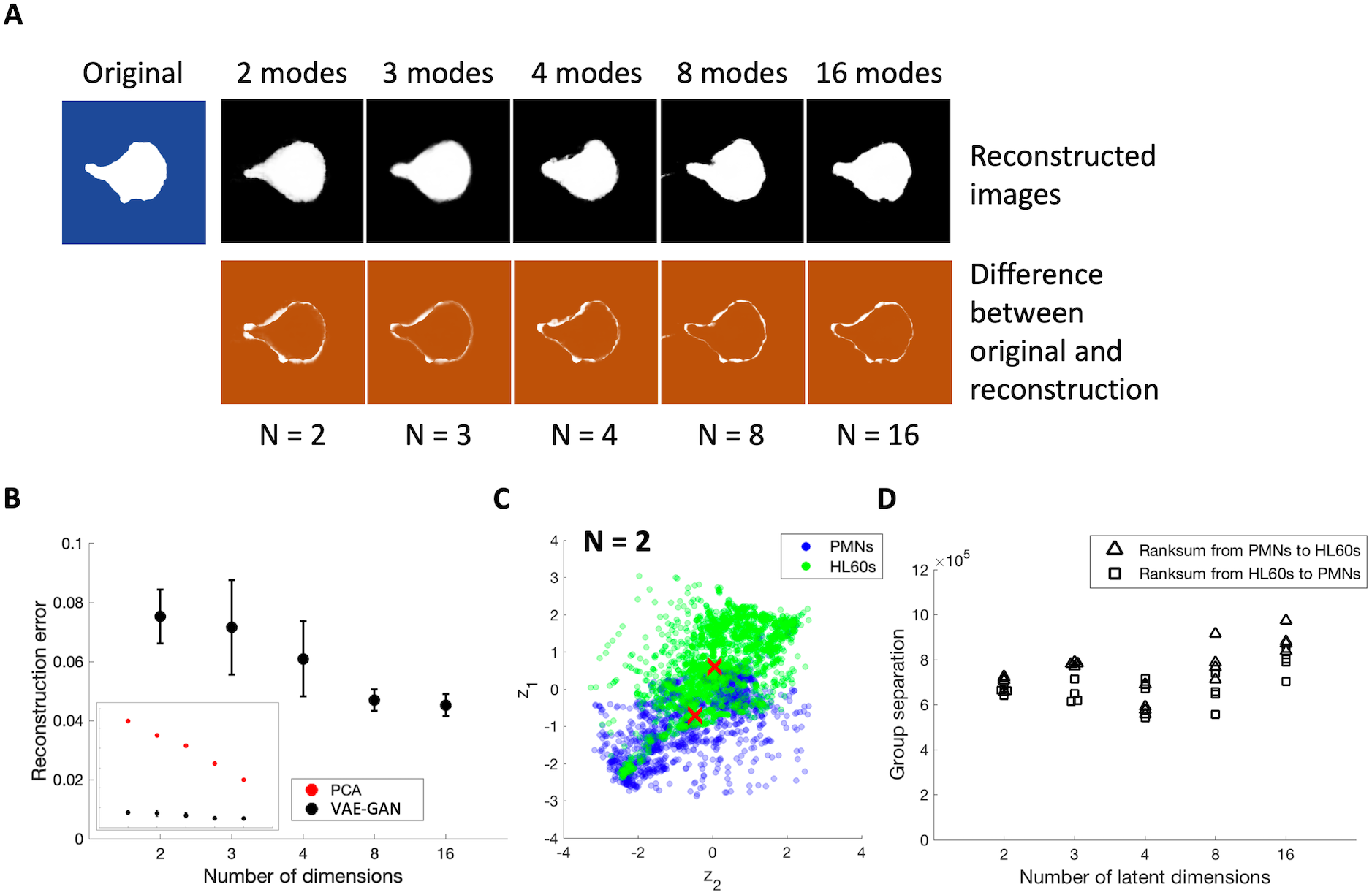
Cell shape reconstruction using a VAE-GAN with different numbers of latent space dimensions. (A) Cell shape reconstruction of a single cell using 2 to 16 latent space dimensions (N = 2 to 16). Original binary mask is shown in blue, reconstructions in black, and difference maps comparing the reconstructed images to the original binary mask in orange. Same original cell as Fig. 2A. (B) Average VAE-GAN reconstruction error for all cells from N = 2 to 16. Error bars show standard deviations from four independent trainings of VAE-GAN for each listed number of dimensions. (C) Scatter plot showing group separation between PMN and HL60 cells in the two latent space dimensions for N = 2 (red crosses = median points for each group) (D) Measure of VAE-GAN group separation ability from N = 2 to 16. Each point represents an independently trained VAE-GAN.

Next, we evaluated the ability of the VAE-GAN to separate the two cell types within the latent space (Fig. 3C-D). Using the same rank-sum metric we had used before (Fig. 2D), we found that the group separation ability between PCA and the VAE-GAN was comparable.

### VAE-GAN latent space encodes information that is consistent with PCA shape space

Prior cell shape studies involving migrating keratocytes (12,13,35) as well as the present work on motile neutrophils have revealed that the major PCA shape modes are biologically interpretable as correlating with expected types of meaningful cell-to-cell variability, including variations in cell area, cell aspect ratio, degree of polarization, and direction of migration (Fig. 1B-D). Given the expected potential relationship between PCA and the VAE-GAN (17,20) we next asked whether the latent space of the VAE-GAN might also contain biologically interpretable information, and whether this information might relate to the shape modes extracted from the PCA shape space.

Using the types of shape variations displayed by the major PCA shape modes as a starting point, we tested whether the same information is also captured in the VAE-GAN latent space. By reconstructing nine cell shapes along each PCA-based shape mode at regular intervals, based on their PCA scores along each mode (from −2SD to +2SD, at 0.5SD intervals), we fed the set of reconstructed shapes into our trained VAE-GAN and determined where each synthetic shape was located within the latent space (Fig. S2A).

Interestingly, all the reconstructed shapes from both shape modes 1 and 2 mapped onto the latent space of N = 2 in the same order as they appeared along their respective shape mode axes, and the two shape modes also appeared to be roughly orthogonal to each other within the latent space, at least near the origin (Fig. 4A). On the other hand, the remaining shape modes (modes 3 and beyond) did not map in the same ordered manner (Fig. S2B). For latent spaces with N = 3 and N = 4, shape modes 1 and 2 mapped even more monotonically and evenly along the entire space, as did shape modes 3 and 4 but to a lesser extent (Fig. 4B, S2C). For N = 8 and N = 16, the ordering of the mapping was further improved, with more and more shape modes mapping evenly across the space (shape modes 1 through 6 shown) (Fig. 4C). In general, these observations suggest that for a given number of latent space dimensions, it is possible to map a similar number of PCA shape modes onto that latent space.

**Figure 4.**
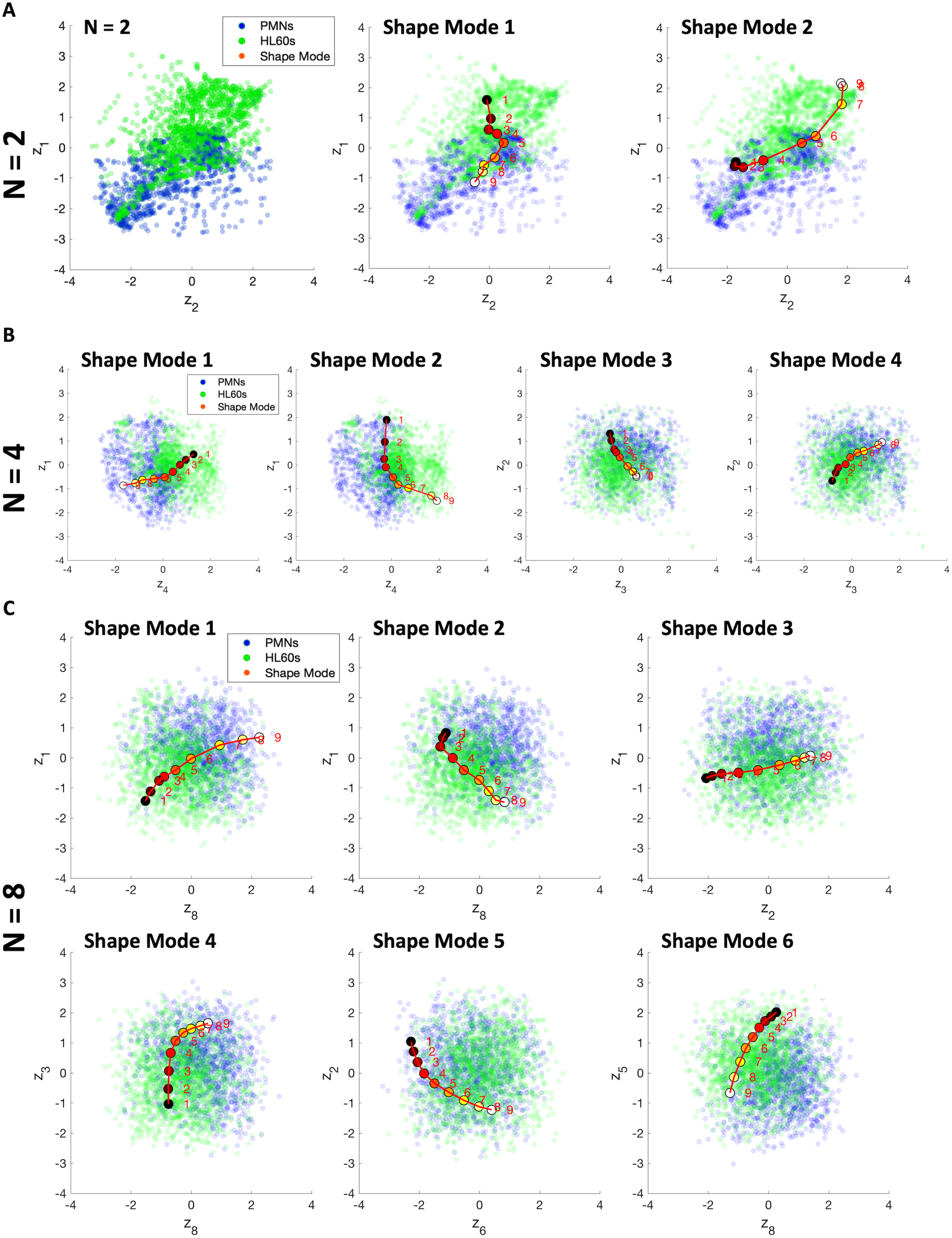
Mapping PCA shape modes to latent spaces. (A) Mapping of shape modes 1 & 2 to latent space of N = 2. Left, scatter plot showing embedding of all points into the two latent space dimensions. Blue = PMNs, green = HL60s. Middle, mapping of principal shape mode 1 onto the same scatter plot. Point 5 represents the average cell shape as determined by PCA and other points represent ordered, evenly spaced synthetic cell shapes sampled along the shape mode 1 axis (see Fig. S2A). Right, mapping of principal shape mode 2 onto the same scatter plot. (B) Mapping of shape modes 1 to 4 to latent space of N = 4. (C) Mapping of shape modes 1 – 6 for latent space of N = 8.

In summary, we have shown empirically that even with the additional non-linear transformation capability of the VAE-GAN, the trained latent space encoded biologically interpretable information that was consistent with the corresponding shape modes we observed in PCA. Specifically, the PCA shape space can be mapped uniquely and monotonically onto the latent space for N = 2, 3, 4, 8 and 16. Combined with our earlier observation that shape reconstruction is much more efficiently encoded in the latent space than in the PCA shape space, this suggests that the optimal decomposition of cell morphology within this data set is a non-linear and non-orthogonal transformation of the biologically interpretable principal components of shape variation as found by PCA.

### Latent space embeddings generated by VAE-GANs are reproducible

Unlike PCA, which is a completely deterministic linear transformation powered by the singular value decomposition of the covariance matrix of the data, VAE-GANs are trained using an iterative optimization methodology where multiple sets of network parameters are initialized randomly and optimized through a machine learning process that involves back-propagation and gradient descent (23,24). Therefore, it is possible that training sessions with different initial conditions may not lead to convergence to identical solutions (33). Specifically, the product of a VAE-GAN is the embedding of the data points within its latent space. Each point represents the encoding of a single input image, and the relative distribution of those points reflects what the network has learned about the relationships among all the images within the data set. In order to evaluate the ability of a VAE-GAN to provide consistent solutions from independent training sessions, we compared the similarities and dissimilarities of latent space embeddings that were trained independently using the same data set but with different initial conditions (random seeds) and different training image input sequences.

For each latent space dimension we explored (Fig. 3), we trained four independent VAE-GANs using the training set of our combined cell shapes. Fig. 5A shows the latent space embeddings of each independent training session for N = 2. For every embedding we can see comparable group separation between PMNs and HL60s. To compare the independent latent spaces to one another, we first used Procrustes analysis to quantify dissimilarities of the embeddings between any two training sessions (36). Procrustes measures the distance, or dissimilarity, between two sets of data points by attempting to optimally conform each set to the other through the use of linear transformations, i.e. operations that include translation, reflection, rotation, and uniform scaling (Fig. 5B). A Procrustes dissimilarity of 0 indicates that the two sets of data points are equivalent after transformation, while a dissimilarity of 1 indicates the opposite.

**Figure 5.**
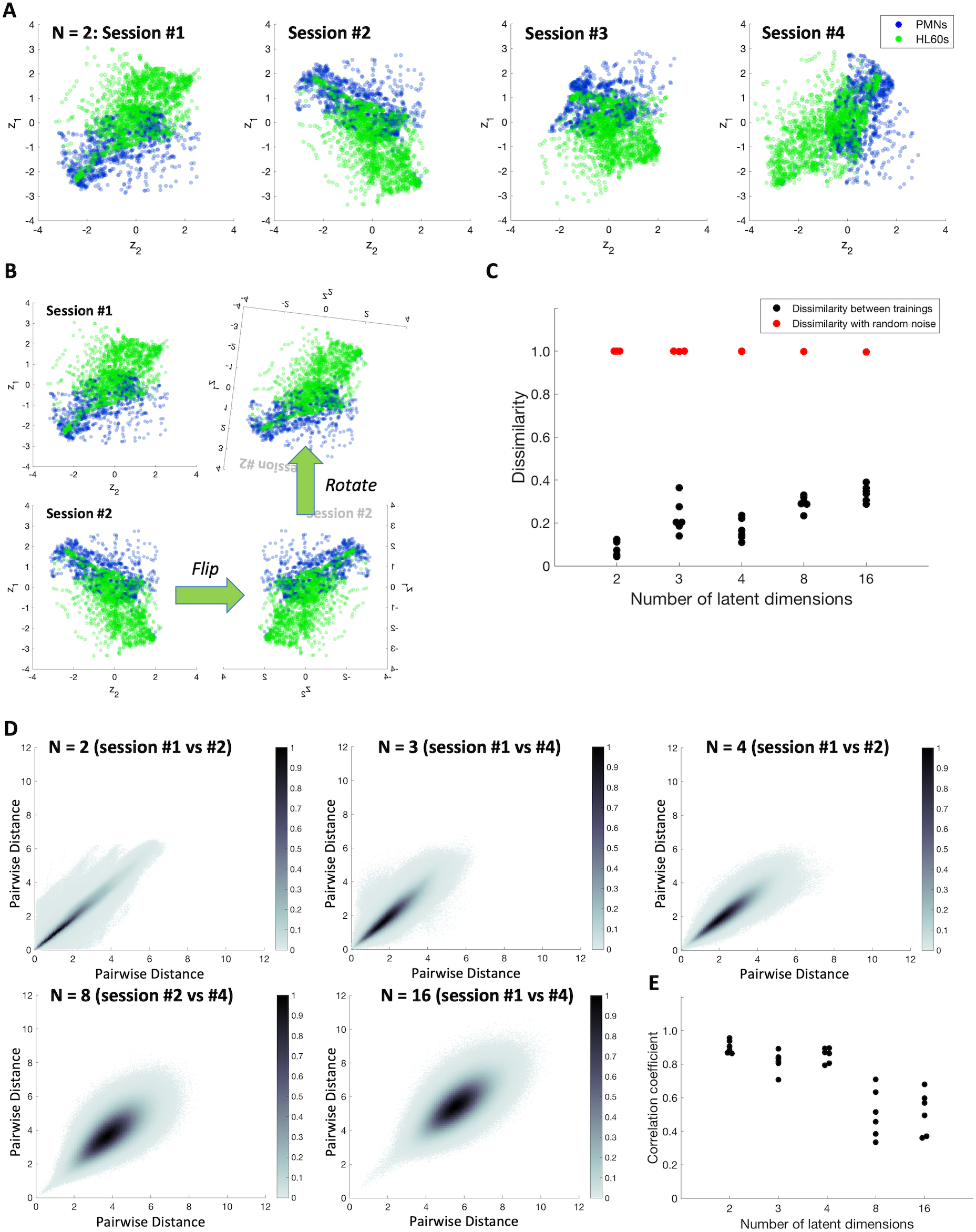
Similarity between independently trained latent spaces. (A) Scatter plots showing embedding of all cells into latent spaces from 4 independently trained networks for N = 2. Blue = PMNs, green = HL60s. (B) Transformations used by Procrustes to compare two sets of data points. (C) Procrustes dissimilarities between independently trained latent spaces using random noise as control for N = 2 to N = 16. Each point represents a pairwise comparison between any two independently trained latent spaces for each indicated dimension. (D) Correlation plots of pairwise distances between latent spaces from two randomly chosen independent training sessions for each indicated dimension. (E) Correlation coefficients between pairwise distances of independently trained latent space for all training pairs across each specified dimension.

We performed Procrustes analysis between all possible pairs of training sessions for each latent space of each dimension for N = 2, 3, 4, 8 and 16, as well as with normally distributed random noise (Fig. 5C). When compared to random noise, all latent space embeddings, regardless of the number of latent space dimensions, resulted in dissimilarity values that were very close to 1, thus indicating that the distribution of points within all latent spaces were very different from random noise. On the contrary, when the embeddings between different training pairs were compared, the dissimilarity values were much lower, ranging from less than 0.13 for N = 2 to no more than 0.4 for N = 16, with an increasing trend of dissimilarities that can be observed as N increases from 2 to 16. This indicates that latent spaces between the individual training sessions were much more similar to each other than to random noise.

To confirm this finding, we applied an alternative strategy by first calculating the pairwise distances between all the points within each latent space, followed by a correlation analysis for the pairwise distances of the latent spaces from each possible pair of training sessions. If two points were close in distance in a particular latent space and were also close in another independent latent space, their pairwise distances between these two latent spaces should be highly correlated. Fig. 5D shows several sample heat-mapped scatter plots of the pairwise distances between two training sessions, randomly chosen from the 4 independent trainings for N = 2, 3, 4, 8 and 16. We found that, in general, the pairwise distances between any two training sessions were highly correlated. In addition, we observed the same trend as in our Procrustes analysis that training sessions with a higher number of latent dimensions were less correlated to each other than those with a lower number of latent dimensions (Fig. 5E). That is, as the number of latent space dimensions increased, the average correlation coefficient between each of the different training pairs decreased, and the spread of correlation coefficients between the different training pairs also increased. Intuitively this result makes sense because VAE-GANs with smaller numbers of dimensions in the latent space are effectively attempting to encode as much information as possible in those few dimensions, and there may be fewer possible ways of doing this efficiently in a small number of latent space dimensions as compared to a larger number with more options.

Therefore, despite random initial network conditions and the stochastic nature of the iterative training process, we have shown that the solutions produced by a VAE-GAN between different training sessions are highly correlated and thus can uncover trends within the data set consistently, as in PCA.

### Cell area is encoded more efficiently in PCA than in VAE-GAN

Having now established the reproducibility of latent spaces within the VAE-GAN, and the monotonic mapping of the PCA shape space onto the latent space, we next turned to the question of biological interpretability. As described above, the top-ranked principal shape modes defined by PCA correspond roughly to a cell biologist’s expectation about meaningful sources of biological variation from one cell to another, as has typically been found in prior applications of PCA dimensionality reduction to cell shape data sets (12,15), and indeed also in applications of PCA dimensionality reduction to other very different kinds of biological data including *C. elegans* posture variation (37) and human facial variation (38). Because the separate dimensions in the VAE-GAN latent space have no direct physical meaning and are not arranged in any meaningful order, interpreting the biological significance of positional variation within the latent space is much less straightforward. Furthermore, the non-linear transformations performed by the encoder and decoder in the VAE-GAN make it challenging for humans to interpret the biological significance, if any, of the “choices” made by the trained neural network in efficiently reducing the dimensionality of the data. These properties have given rise to the general opinion that deep neural networks are “black boxes” that can perform well at classification and prediction tasks but fundamentally cannot be used to gain mechanistic insight into the underlying causes of variation in a data set (39).

We set out to explore this issue by mapping two biologically meaningful and directly observable metrics for our crawling human neutrophils onto both the PCA shape space and the VAE-GAN latent space, namely cell area and cell speed. For neutrophils crawling on glass coverslips underneath an agarose overlay, each individual cell maintains a very constant projected cell area over time (28), but individual cells may be quite different in size from one another. As noted above, the first principal shape mode derived from PCA appears to the human eye to roughly correlate with cell area, and of course it makes sense that this basic geometrical property of cell shape must be encoded in the cell shape data set used in these analyses. Cell speed, on the other hand, can vary from moment to moment for an individual cell (28) and is not directly or explicitly encoded in the shape data alone, although correlations between shape and speed may exist and indeed have been demonstrated in other motile cell types (12,14)

To examine the encoding of cell area within the PCA shape space more rigorously, we calculated the area of each cell shape in the data set by finding the area of its corresponding binary mask. By plotting cell area against the PC scores of different shape modes we found that shape mode 1 is indeed more closely correlated to cell area as compared to the other modes (Fig. 6A-C). Next, we modeled this relationship by constructing a set of linear regression models that utilized the scores from the first 2, 3, 4, 8, and 16 shape modes to predict the cell areas of our cell shape test set (n = 159). We were able to demonstrate the predictive abilities of these models by plotting the predicted cell areas against the actual cell areas (Fig. 6D, S3A). By calculating the root mean square (RMS) errors and correlation coefficients between the predicted and actual cell areas for the test set, we evaluated the prediction accuracy of our linear regression models, and we observed that the predicted results for all our models (from 2 to 16 shape modes) were highly correlated with the ground truth (actual cell areas), with correlation coefficients that ranged from 0.97 (when using only the top 2 shape modes), to 0.99 (when using the top 16 shape modes) (Fig. 6E-F). As expected, the prediction accuracy was very high even when we used only the top 2 shape modes for our model, with the regression coefficient of shape mode 1 contributing the most to the predicted areas in each model (Fig. 6D, bottom). This is consistent with our earlier qualitative interpretation of shape mode 1 as being the major descriptor of cell area variations.

**Figure 6.**
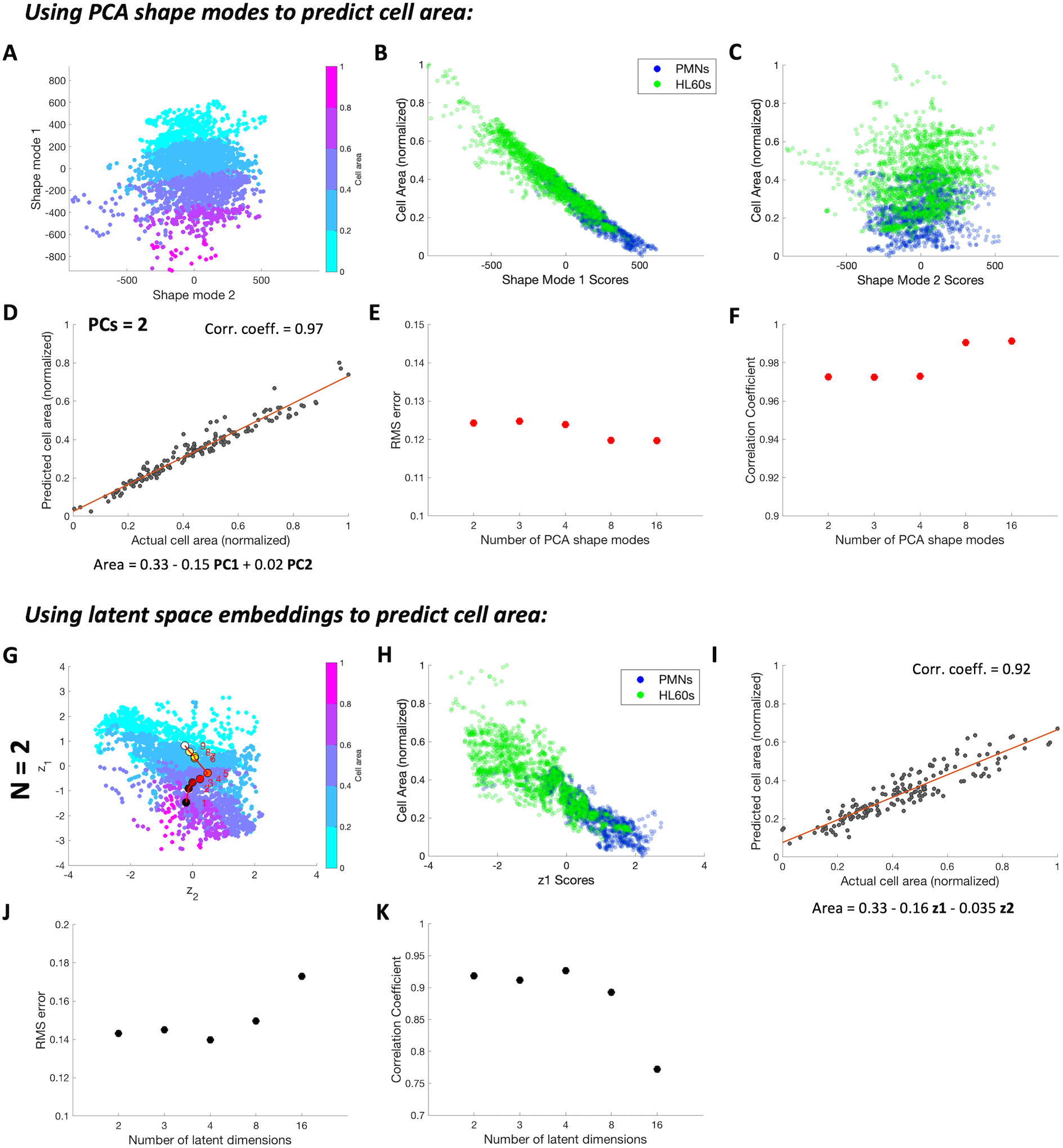
Predicting cell area using PCA shape modes and VAE-GAN latent spaces. (A) Scatter plot of PC scores for principal shape mode 1 vs. principal shape mode 2, color-coded by cell area. (B & C) Cell area versus PC scores for principal shape modes 1 & 2. Blue = PMNs, green = HL60s. (D) Predicted versus actual cell area using the first two shape modes in a linear regression model. Equation shows regression intercept and coefficients for the two shape modes used. (E & F) Average root mean square (RMS) error and correlation coefficients of cell area prediction accuracy for linear regression models using 2 to 16 principal shape modes (G) Scatter plot of latent space embeddings for z_1_ versus z_2_ for N = 2, color-coded by cell area and overlaid with PCA shape mode 1 (red). (H) Cell area versus z_1_ embeddings for N = 2 (I) Predicted versus actual cell area using z_1_ and z_2_ for N = 2 in a linear regression model. Equation shows regression intercept and coefficients for z_1_ and z_2_. (J & K) Average RMS error and correlation coefficients of cell area prediction accuracy for linear regression models from N = 2 to 16.

We then extended our analysis and to ask whether cell area variations are also consistently represented in the latent spaces of our VAE-GANs, when trained on the same data. From Fig. 4 we observed that shape mode 1 could be consistently mapped across the latent space; correlation between cell area and latent space embeddings could also be observed along a subset of latent space dimensions (Fig. 6G-H, S4A-B). By building a set of linear regression models using latent spaces with 2 to 16 dimensions, we evaluated their abilities for predicting cell area for the same test set we used in our PCA approach (Fig. 6I). As expected, these models also displayed reasonably high predictive accuracy for cell area, similar to our PCA-based models, with correlation coefficients between the predicted and actual cell areas ranging from 0.92 to 0.89 from N = 2 to N = 8, while decreasing slightly to 0.72 for N = 16 (Fig. 6J-K, S4C-D). Notably, though, the prediction accuracy for cell area was lower for VAE-GANs of all dimensions as compared to PCA, and the accuracy actually decreased for higher N in the VAE-GAN, in contrast to the increase for higher N for PCA (Fig. 6F, J).

In brief, the fundamental cell geometrical property of area is indeed encoded within the latent space in a simple way that can be captured with a linear regression model, but this encoding is relatively less efficient than the equivalent encoding in PCA and degrades for latent spaces with high dimensionality.

### Neutrophil cell speed is not represented in PCA or VAE-GAN through cell shape

Variations in cell area serve as an important measure in understanding the morphological heterogeneity of a cell population, and were therefore useful to us as a positive control for latent space interpretability. Whereas cell area by nature must be fully encoded in our shape representations of cells, there is no such guarantee that cell speed, another fundamental and biologically important quantitative descriptor of motile cells, would necessarily be directly encoded by shape. Based on prior results in other motile cell types including fish epidermal keratocytes (12) and *Dictyostelium discoideum* (14) we initially hypothesized that cell speed may be embedded implicitly within these shape representations, specifically as a combination of shape modes (in PCA) or latent space dimensions (in VAE-GANs). To explore that possibility, we adopted the linear regression-based methodology used in our cell area analysis above.

For each single-cell time-lapse movie in our data set, we first calculated the migration speed of the cell between two successive time points (a time segment) by dividing the displacement of the cell’s centroid between the two time points with the imaging time interval. We then approximated the instantaneous speed of the cell at each time point by averaging the cell’s calculated speed from the time segments before and after that time point. When we plotted cell speed at each time point against the PC scores of the corresponding cell shapes, we observed no obvious trends correlating cell speed with PC scores from any of the major shape modes (Fig. 7A-C, S5A). This observation was further confirmed by the construction of a set of linear regression models that attempted to utilize the PC scores from the first 2, 3, 4, 8, and 16 shape modes to predict the cell speed of our cell shape test set (n = 159). We evaluated the predictive abilities of these models by plotting predicted cell speed against actual (measured) cell speed (Fig. 7D-E, S5B), and found that the correlation between predicted and actual cell speed is low, with correlation coefficients ranging from 0 (when using 2 shape modes) to 0.41 (when using 16 shape modes) (Fig. 7F-G), which can be interpreted as having weak correlations at best, indicating that these regression models based on PCA shape modes for this data set have little predictive power in predicting cell speed using cell shapes alone.

**Figure 7.**
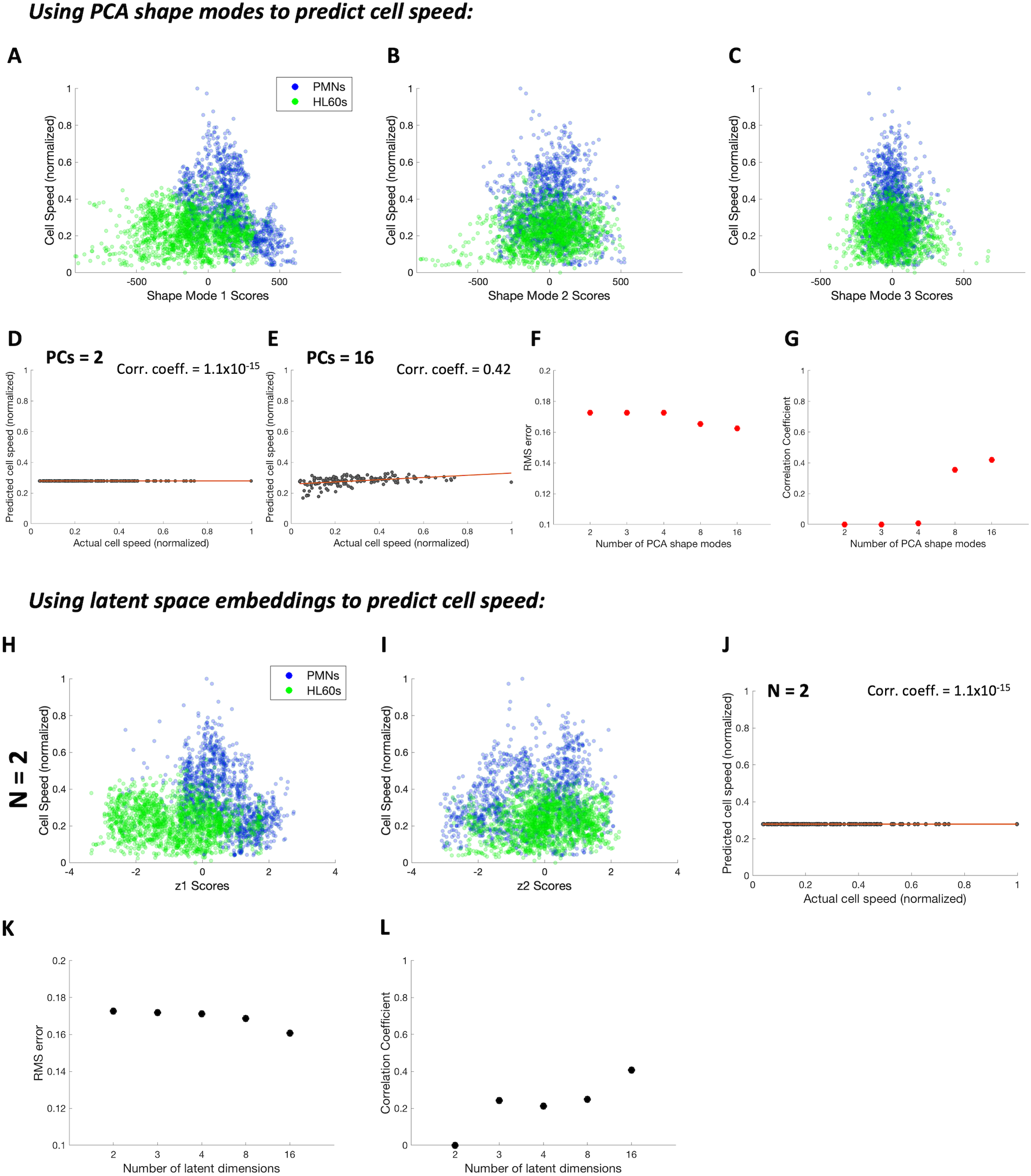
Predicting cell speed using PCA shape modes and VAE-GAN latent spaces. (A – C) Scatter plots showing cell speed versus PC scores of principal shape modes 1 to 3. Blue = PMNs, green = HL60s. (D & E) Predicted versus actual cell speed using the first 2 and 16 shape modes in linear regression models, respectively. (F & G) Average RMS error and correlation coefficients of cell speed prediction accuracy for linear regression models using 2 to 16 principal shape modes. (H & I) Scatter plots showing cell speed versus z_1_ & z_2_ embeddings for N = 2. (J) Predicted versus actual cell speed using z_1_ and z_2_ for N = 2 in a linear regression model. (K & L) RMS error and correlation coefficients of cell speed prediction accuracy for linear regression models from N = 2 to 16.

Next we asked whether cell speed could be recovered from the latent space embeddings of our trained VAE-GANs. We plotted cell speed against the latent space embeddings for N = 2 to N = 16, and similar to what we found in PCA, we could detect no obvious linear correlation between cell speed and latent space embeddings (Fig. 7H-I, S6A-C). This was quantified by constructing the corresponding linear regression models to use these latent space embeddings in predicting cell speed (Fig. 7J, S6D). Just as in PCA, we observed little to no correlation between predicted versus actual speed, indicating that cell speed was also not represented in the latent spaces of our trained VAE-GANs (Fig. 7K-L).

Notably in both the PCA shape space and the VAE-GAN latent spaces there do appear to be zones of cells that are substantially faster than average, particularly for the primary human cells (PMNs), but these tend to cluster near the origin and are not monotonically correlated with any single latent parameter or principal mode (Fig. 7A-C, H, S5A, S6A-C). In other words, cells with average or common shapes may be moving at any speed, while cells with shapes that are significant outliers are more likely to be slow. While it might be possible to achieve better predictive value for cell speed with an ad hoc nonlinear regression, this seems unlikely to yield meaningful mechanistic insight.

### VAE-GANs can integrate cell speed with cell shapes in the latent space

Next, we turned to the question of whether the architecture of the VAE-GAN could be exploited to encode speed into the latent space directly. The encoder in our VAE-GAN was originally designed to extract features from multi-channel 2D images using deep convolutional layers (24). Previously we did not take advantage of this feature, as our binary masks were encoded in a single channel. This gave us the opportunity to attempt to incorporate speed encoded as an additional image channel, along with cell shape (Fig. 8A). The additional speed channel took the form of a centered square with a uniform pixel intensity value that reflected cell speed, with pixels outside of the square having intensity values of zero (Fig. 8A, right). The normalized speed of a cell was associated with its shape by being represented as the uniform pixel intensity value within the speed square, where a cell that is moving fast at a particular time point would have an intensity value higher than that of a slower moving cell. The area of this speed square was identical across all cell shapes. So that the training of the network would balance shape and speed information relatively equally, we chose a size for the speed square based on the average number of pixels that varied among all the binary masks within the entire cell shape data set (see *Methods*). The VAE-GAN was then trained again with the same data set of cell shapes we used earlier, along with their corresponding speeds encoded in the previously unused green channel.

**Figure 8.**
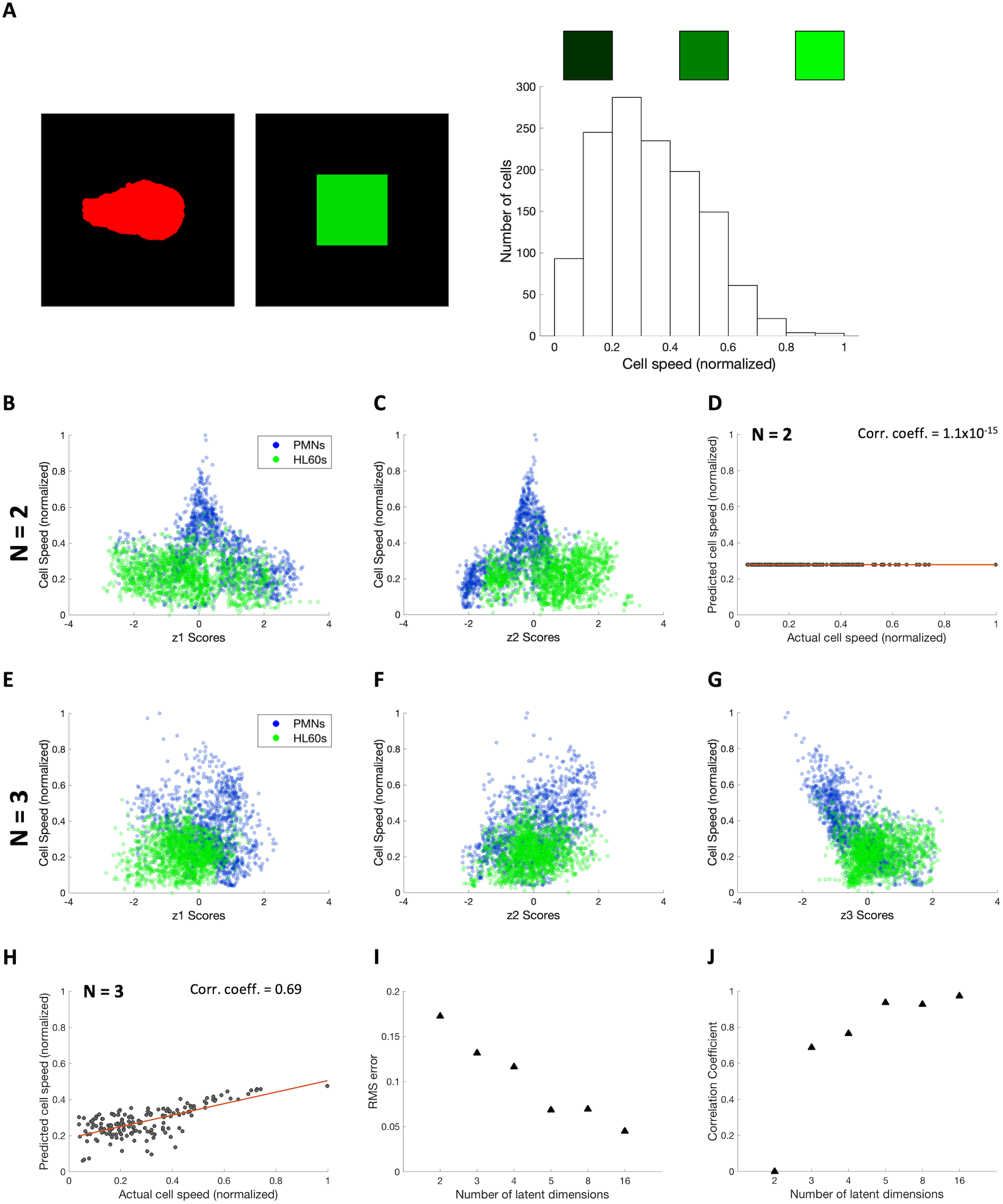
Incorporating cell speed in cell shape analysis using a VAE-GAN. (A) Left, cell shape (red) and cell speed (green) encoded in two separate image channels as training data for a VAE-GAN. Right, histogram of cell speeds, showing green intensity coding of speed. (B & C) Scatter plots of cell speed versus z_1_ & z_2_ embeddings for N = 2. (D) Predicted versus actual cell speed using z_1_ and z_2_ for N = 2 in a linear regression model. (E - G) Scatter plots of cell speed versus z_1_ to z_3_ embeddings for N = 3. (H) Predicted versus actual cell speed using z_1_ to z_3_ for N = 3 in a linear regression model. (I & J) Average RMS error and correlation coefficients of cell speed prediction accuracy for linear regression models from N = 2 to 16

Using this image-based speed encoding scheme, we re-trained the VAE-GANs using latent spaces that contained 2, 3, 4, 5, 8, and 16 dimensions. We then plotted cell speed against different dimensions of the latent space embeddings, and observed that with N = 2, there was no obvious correlation between latent space embedding and speed (Fig. 8B-C, S7A). This was confirmed by using the latent space embeddings to predict cell speed through linear regression, where the correlation coefficient between the predicted and actual speed was calculated to be close to 0 (Fig. 8D). From N = 3 onwards, however, we began to observe latent space dimensions that appeared to be correlated with speed (Fig. 8E-G, S7B). This was supported by our linear regression models for speed prediction (Fig. 8H, S7C), where the correlation coefficients between predicted and actual speed ranged from 0.69 to 0.97 for N = 3 to 16, indicating that the additional speed information has been successfully incorporated into the latent space embeddings in a simple, monotonic way that can be recovered by the linear regression models (Fig. 8I-J).

By utilizing the latent spaces that have been trained to associate cell speed and shape, we can now make inquiries about how those two cell features are related. The first question we asked was whether we can distinguish a fast-moving cell versus slower-moving cells based on shape. We answered this question by first dividing all the cell shapes within our data set into bins based on speed and calculating the median of the corresponding latent space embeddings for each bin. We then located these embeddings, representing each with the actual shape of a migrating cell that was close to the median for each bin (Fig. 9). We applied this process to trained latent spaces with dimensions N = 2, 3, 5, 8, and 16, and observed that the fastest moving cells were homogeneous in shape (Fig. 9, right column), while the slower moving cells had heterogeneous shapes (Fig. 9, left column), with cells migrating at intermediate speeds exhibiting shapes that were in between (Fig. 9, middle column). This suggests that, while the fastest moving cells all tended to be polarized in shape migrating forward, the slower moving cells could have any shape. Notably the average cell shapes for both PMNs and HL60s in this particular data set were fairly polarized (Fig. 1A, right). This explains why the fastest-moving cells tended to be embedded close to the origin in both the PCA shape space and the VAE-GAN latent spaces.

**Figure 9.**
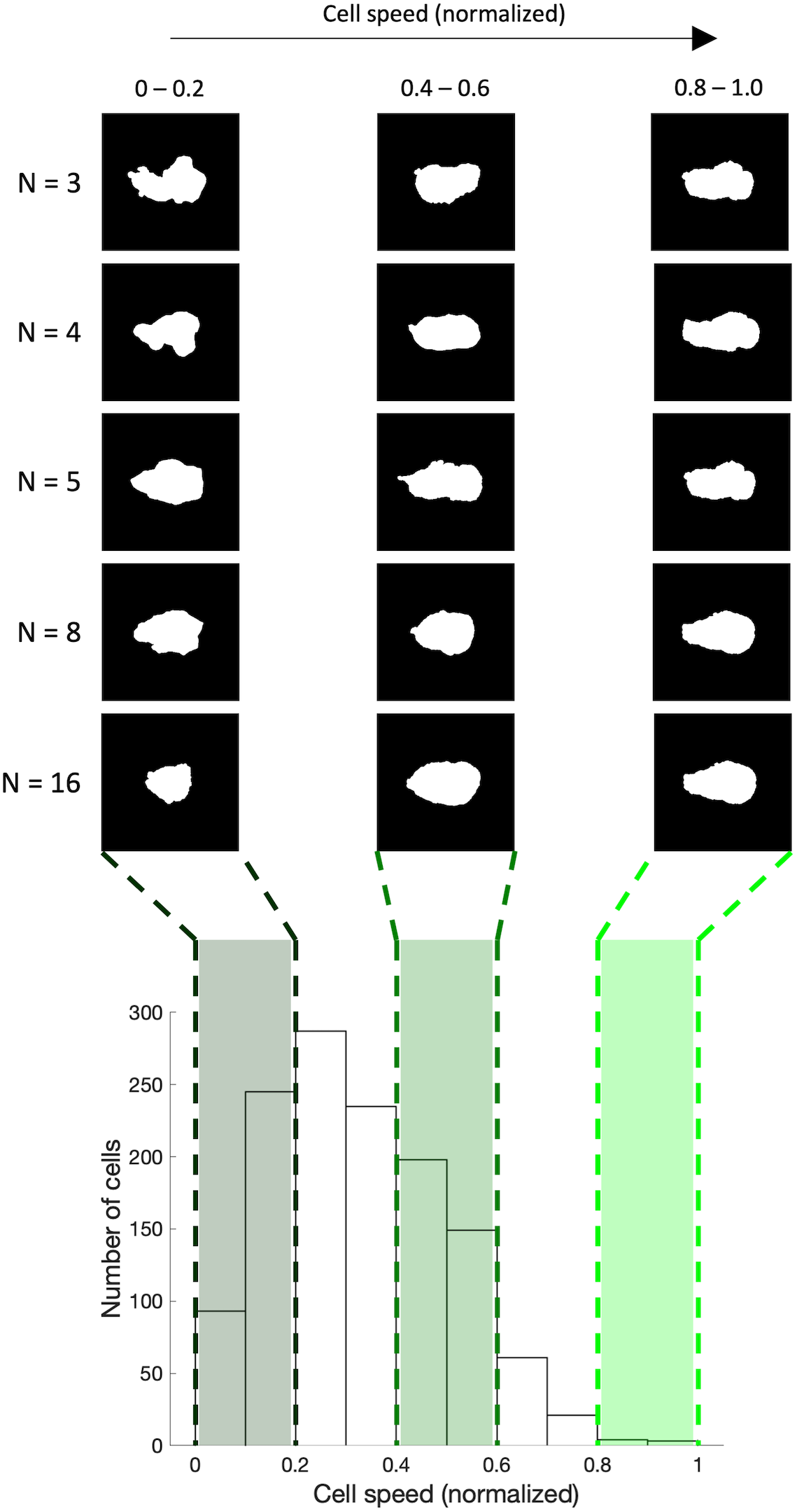
Relating cell speed to cell shape. Top, median cell shapes of slow (0 – 0.2), medium (0.4 – 0.6), and fast (0.8 – 1.0) speed bins for N = 3 to 16. Bottom, histogram of all cell speeds with slow, medium and fast bins indicated.

## DISCUSSION

The complex shape of an amoeboid-like cell can be conceptualized as a readout of the intracellular and extracellular forces acting on the cell membrane (11). In addition, the expected coupling between a cell’s shape or structural organization and its function suggests that changes in cell morphology of migrating cells can be leveraged to enhance our mechanistic understanding of a cell’s motility behavior (28). In this paper we developed a quantitative framework that compactly represents the cell shape of migrating human neutrophils (PMNs and HL60s) in low-dimensional space. By extracting the major modes of shape variation and integrating experimentally measured motility parameters into predictive models, we can now make inquiries into the connection between cell shapes and motility behavior.

Previous studies on connections between cell shape and motility in other cell types have demonstrated that cell aspect ratio can be strongly correlated with speed. In fish epidermal keratocytes, rapidly-moving cells are oriented with their long axis perpendicular to the direction of motion, and this widening of the leading edge is driven by the force balance between actin polymerization and tension in the plasma membrane (12,35). In the cellular slime mold *Dictyostelium discoideum*, rapidly-moving cells are oriented with their long axis parallel to the direction of migration, and this elongation appears to reflect their degree of cytoskeletal polarization (14). For neutrophils, we had initially expected that front-rear polarization (as captured in the PCA-based shape modes 2 and 4) would correlate with speed, but our measurements here do not support that simple hypothesis. However, the fact that we can jointly embed both speed and shape information into a single latent space offers the possibility that the relationship between neutrophil cell shape and speed is predictable in some way, although more work will be required to understand exactly how. A very promising avenue will be to include fluorescent protein distribution images for cell structural components known to be critical for determination of both cell shape and cell speed, such as actin filaments and non-muscle myosin II (28).

In addition to our interest in building a quantitative framework for exploring the connections between cell shape and cell movement behavior in neutrophils specifically, more generally in this work we aimed to compare directly both traditional (PCA) and modern (VAE-GAN) approaches to dimensionality reduction for image-based cell biological data sets. Along with prior work on cell shape specifically (9,12,14,15), dimensionality reduction for complex image data has proved to be very useful for characterization of animal behavior phenotypes. For example, simple PCA characterizing the posture of crawling nematodes (*Caenorhabditis elegans*) in videomicroscopy images was sufficient to establish that the first two principal modes of posture variation (“eigenworms”) reflect an oscillatory circuit for forward crawling, and the third mode represents occasional “U-turns” when worms change direction (37). Importantly, these same principal modes, derived strictly from observations of wild-type animals, could be used for a compact quantitative description of phenotypic variation in behavior of several hundred mutant strains of *C. elegans* with behavioral defects (40). For more complex animals such as the fruit fly *Drosophila melanogaster*, PCA alone is not sufficient to capture postural variations reflecting movement behavior in an interpretable way, but non-linear dimensionality reduction methods on image data for freely moving fruit flies have been able to reveal stereotyped patterns of complex behaviors (41,42).

Recent advances in the design of deep convolutional neural networks (CNN) such as the GAN (23), and ongoing rapid improvement in computational power, particularly the adoption of GPUs for biological computing (43), have led to a recent surge of interest in the possibilities of deep learning for cell biology. To date, nearly all cell biological applications of this technology have used some form of supervised learning for classification tasks (44). For example, CNNs have been effectively used for cell-by-cell classification with goals including predicting differentiation of stem cells (45), classifying cell cycle status (46), and identifying cell types based on their motility behavior (47). Pixel-by-pixel classification strategies for cell images have also been fruitfully applied to the problem of image segmentation, using human-annotated “ground truth” images for training purposes (48). A different form of supervised learning enables prediction of one image type from another, such as predicting a high-resolution image from a low-resolution input (49,50), or predicting a fluorescence image of a particular labeled structure from a transmitted light image (51,52). Unsupervised deep learning, in contrast, has not been widely adopted.

Dimensionality reduction is a natural application for unsupervised deep learning (17). While traditional dimensionality reduction methods such as PCA are very easy to implement and understand, they are severely limited by the constraints that all transformations must be linear and all independent dimensions must be orthogonal to one another by construction. There is no a priori reason to expect that the most meaningful compact representation of cell shape, or indeed any cell structural feature, should intrinsically obey these mathematical rules. Indeed, for our neutrophil cell shape data set, we found that a VAE-GAN constrained to encode as much information as possible in just a handful of dimensions (N = 2, 3, 4, etc.) could perform substantially better in accurate image reconstruction than a PCA-based decomposition of much higher dimensionality (up to 512). This demonstrates that the most efficient encoding of cell shape is not, in fact, linear or orthogonal in nature.

Although we were able to demonstrate that independent trainings of VAE-GANs generate reasonably reproducible latent spaces, particularly when constrained to a very small number of dimensions, direct biological interpretation of the latent space dimensions remains challenging. It is encouraging that the well-defined PCA shape space for this data set maps smoothly and monotonically onto the VAE-GAN latent spaces, but this approach will not be able to be used directly for interpreting latent spaces generated from more complex types of training where an independent PCA-based (or other linear) analysis is not feasible. The fundamental problem is that the separate dimensions within the VAE-GAN latent space, while constrained in their number, have no intrinsic physical meaning, and are not sorted into any rational order based on importance or significance in the way that principal modes from PCA are sorted automatically by the amount of variance explained. To this end, other VAE architectures may be more promising than the VAE-GAN. In particular, the ß-VAE architecture uses the penalty calculation in the training process to favor sparse representation in the latent space as well as reconstruction accuracy (53). Because of this, it is possible to sort the latent space dimensions after training with respect to the amount of variance explained by each dimension, highly analogous to the sorting of dimensions in PCA (54). An alternative approach would be to use archetype analysis autoencoders such as AAnet (55). These seek to reduce dimensionality in complex data sets by identifying a minimal number of extreme “pure types” or archetypes in the data space and describing other samples as convex combinations of these archetypes. Both ß-VAE and AAnet might be expected to generate latent spaces that are more easily directly interpretable than VAE-GAN.

Historically, the field of quantitative biology has benefited from the use of experiments and theory iteratively to inform one another, in order to refine our conceptual and mathematical models of the biophysical rules governing a cell’s state or function. Our work here has shown that unsupervised deep learning can be harnessed as a powerful complementary tool that can extract interpretable biologically relevant variations from image data and help further our understanding in the mechanisms of cellular function. The great power and flexibility of using deep CNN for embedding multiple distinct kinds of data in the latent space is particularly promising. Here we have demonstrated that cell speed information can be compactly embedded along with cell shape, and in the future it will be particularly interesting to also incorporate fluorescence images showing the spatial distribution of particular proteins of interest known to be important for cell motility.

Our results are significant in demonstrating the utility of unsupervised, as compared to supervised, deep learning approaches for cell biological discovery. Unsupervised learning does not require prior labeling of the data, nor the intrinsic property of having known discrete classes in the data. However, concerns about the “black-box” nature of the latent space have limited the popularity of unsupervised learning methods in scientific inquiries that aim to extract biologically interpretable information. In our work, we used PCA to guide our exploration of the latent space, allowing us to gain insight into neutrophil motility behavior for a data set that contains continuous shape variations with no discrete classes. Attempting to shine light into the black box of the VAE-GAN, we have seen a latent space that is not entirely unfamiliar, but instead can be rationally mapped with clearly interpretable variation as measured by a deterministic and linear dimensionality reduction technique (in this case, PCA). The great potential power of CNNs for aiding in the interpretation of large and highly complex biological data sets indicates that further efforts along these lines will be well-justified.

## ACKNOWLEDGMENTS

We would like to express special gratitude to Gregory Johnson and Rory Donovan-Maiye of the Allen Institute for Cell Science for their advice and guidance on the training of the VAE-GAN used in this paper, which is based on their work on the integrated model of the cell (24).

We acknowledge all members of the Theriot lab for helpful discussions, and thank Nathan Belliveau and Lorenzo Labitigan for comments on the manuscript. This work was supported by the Howard Hughes Medical Institute (J.A.T.). C. K. C. was supported by the Stanford Cellular and Molecular Biology Training Grant (NIH T32-GM007276). A.H. was a Stanford Bio-X Bowes Fellow, an Onassis Foundation Scholar and was also supported by the Foundation for Education and European Culture.

## MATERIALS AND METHODS

### Culturing of HL60 cells

Neutrophil-like HL60 cells were maintained in RPMI media (Gibco 22400), supplemented with 10% heat-inactivated fetal bovine serum (hiFBS) (Gemini 900-108, heated in water bath at 55 ^o^C for 30 minutes), 100 U/ml penicillin, 100 μg/ml streptomycin and 0.25 μg/ml amphotericin B (Gibco 15240). The cultured cells were incubated at 37 ^o^C in 5% CO2 and diluted once every 2-3 days into a density of 2×10^5^ cells/ml. To differentiate the HL60s, cells were diluted in RPMI media containing 1.3% DMSO (Acros 61097) with an initial density of 2×10^5^ cells/ml. For all experiments, only cells differentiated for 5-6 days were used.

### Preparation of primary human neutrophils

Human polymorphonuclear leukocytes (PMNs) were isolated from blood samples by density gradient separation using PolymorphoPrep (Axis-Shield PoC AS, Norway). Briefly, peripheral blood from two healthy adult volunteers was collected in K2 EDTA-coated tubes (BD Vacutainer, Becton Dickinson). Within 30 minutes of drawing, 5 mL of blood was carefully laid over 5 mL of PolymorphoPrep in a 15 mL conical bottom polyproylene Falcon tube (Fisher Scientific). Tubes were then centrifuged at 650 g for 35 minutes in a swinging bucket rotor with acceleration set to 7 and no brake for deceleration (deceleration set to 0) using a Thermo Fisher Sorvall Legend XT centrifuge. The plasma (top yellow layer) and mononuclear cells (second clear layer) were removed with a 5 ml syringe using a 16 gauge blunt needle, allowing the harvest of PMNs with a 3 ml syringe and a 16 gauge blunt needle. Harvested PMNs were then re-suspended in an equal volume of 0.5x neat RPMI for osmotic equilibration and centrifuged at 350 g for 10 minutes (with acceleration and deceleration set to 9). Upon removal of the supernatant, the PMN pellet was re-suspended in L-15 (21083-027, Gibco) to densities comparable to those used for HL60s. PMNs were maintained at 37 ^o^C and used in experiments within 6 hours after isolation.

### Nuclear staining of cells

At the beginning of each experiment, about 10^5^ cells (differentiated HL60s or PMNs) were spun down at 500 g for 5-10 minutes and resuspended in 1 mL of L-15 media. 1uL of stock 1mg/mL Hoechst 33342 stain was added to the L-15 cell suspension to reach a final concentration of 1ug/mL. The stain-containing cell suspension was then incubated at 37 ^o^C for 20 minutes before being re-pelleted at 500 g for 5-10 minutes and resuspended in 1 mL of L-15 in preparation for being used in the under-agarose assay.

### Under-agarose 2D cell migration assay

In this assay, cells were confined to migrate in a 2D environment between an agarose pad and a fibronectin coated #1.5 glass coverslip. Clean coverslips were first incubated with 10 μg/ml fibronectin in PBS (Sigma F2006) for 1.5 hours prior to experiment. After incubation the coated coverslips were first rinsed with PBS before being stored at 37 ^o^C in complete RPMI media until cell plating (see below).

To cast the agarose pad, we first prepared a 2% (20mg/mL) low melting point agarose (Invitrogen 16520) solution by dissolving the agarose powder in heated neat L-15 media (solution A) that was kept in a 37 ^o^C water bath until use. Next, we prepared a 2x stock solution with neat L-15 media containing 20% hiFBS (solution B). f-MLP (Sigma F3506) was added as chemoattractant to solution B at a final concentration of 2 nM to induce chemokinesis. Solutions A and B were then mixed at a 1:1 ratio (with a final concentration of 1% agarose, 10% hiFBS, and 1nM fMLP), and 1mL of this mixture was pipetted immediately into a 25mm diameter mold to cast the agarose pad. The agarose was kept at room temperature for at least 20 minutes to solidify.

To plate the cells for imaging, the washed Hoechst-stained cell suspension (see above) was first spun down with 500g for 5-10 minutes, and then concentrated to a final volume of 10 μl by removing the supernatant (final cell concentration = 10^4^ cells/μl). The concentrated cell solution was then resuspended and added dropwise to the center of a fibronectin coated coverslip (see above). The solidified agarose pad was immediately overlaid above the 10 μl cell solution. The gel-covered coverslip was transferred to a microscope coverslip holder and used for time-lapse imaging. Typically, this setup was incubated at 37 °C for about 20 minutes until the agarose pad would start to compress the cells. After the cells were fully confined under the pad, the cell area would increase, and the brightness of the halo around the cell would decrease in phase-contrast images. We only started imaging after the cells were fully confined under this definition.

### Microscopy and data acquisition

Cell shapes from human neutrophils migrating on a 2D plane and confined under agarose were obtained from live cell imaging using phase-contrast time-lapse microscopy. To image the shape change of migrating neutrophils with high spatiotemporal resolution, we used a Nikon Eclipse Ti epifluorescent microscope with a 100x oil-immersion objective and a 1K back-thinned EMCCD camera, Andor iXon3 (Andor, Belfast, United Kingdom). Acquisition rate was set to 17-24 frames/min (2.5-3.5 s time intervals) for PMNs and 6-40 frames/min (1.5-10 s time intervals) for HL60s. The higher frame rate used for PMNs was necessary due to their more dynamic shape change during migration, when compared to HL60s. For each frame the cell was imaged via two different channels, one for phase contrast imaging (for the extraction of cell shapes) and one for epifluorescent imaging of the Hoechst-stained nucleus signal (for alignment of the nuclei as the cell center). The size of the raw image acquired was therefore 1024 by 1024 pixels by 2 color channels per frame. Temperature was kept at 37 ^o^C with a hot air heater (AirTherm) and humid by using a custom made sample cover with damp lab wipes. Time-lapse movies of various lengths and time intervals were collected by imaging 17 PMN cells extracted from two healthy adult volunteers, amounting to a total of 1298 cell shapes. Imaging for HL60 cells was performed on 209 different cells, totaling 1715 cell shapes.

### Data preprocessing and extraction of cell shapes

Individual migrating neutrophils from the raw microscopy images were first identified programmatically using custom Matlab code, and both the phase contrast and Hoechst-stained channels were cropped into smaller images of 401 by 401 pixels using the centroid of the nucleus as the image center. Each cropped phase-contrast image was then manually segmented using the magic wand selection tool in Adobe Photoshop and saved as a binary mask. Cell contours from different frames and cells were extracted by feeding the binary masks through Celltool (9) and imported into Matlab as 2 x 300 matrices that contained the xy coordinates of 300 equidistant points along the edge of each cell. Other than down sampling the original cell contour by discretizing it into 300 two-dimensional points, the complete shape of the cell is otherwise preserved with this representation. The contours were then aligned programmatically along the +x direction in Matlab using the direction of cell movement (tracked using the change in nucleus centroid positioning) and by minimizing the mean squared distances between contour points on consecutive frames.

### Principal component analysis and shape mode visualization

The numerical representation of each cell contour (300 points, each with an x and y coordinate) was first serialized and aligned (300 points x 2 coordinate values in the x and y directions) into an M by 600 matrix, where M is the number of contours in the data set, and 600 is the number of dimensions occupied by each contour. PCA was then performed on the resulting matrix in Matlab, which produced a set of orthogonal eigenvectors, ranked in order of their respective variations within the data set, and the PC score of each contour along these vectors. Visualization of the shape modes were done by sampling equidistant points along the eigenvectors that contain the most variations, from −2 to +2 standard deviations. Since each point in this shape space contains 600 dimensions, the coordinates of these equidistant points can be used to reconstruct the cell contour represented by those points.

### Training the VAE-GAN

A total of 3172 cell shapes were segmented into binary masks, including both PMN and HL60 cells. Out of that, the binary masks of 3013 cell shapes (of which 1298 are PMNs and 1715 are HL60s) were randomly selected as the training set to train the VAE-GAN. A wide range of values for each hyperparameter was tested, and the final values were selected to ensure a quick and stable convergence of the reconstruction error.

### Test set selection

The remaining 159 images (about 5% of the total number of segmented images) were used as a test set to evaluate whether overfitting had occurred. The same set of test images were also used throughout the various PCA versus VAE-GAN comparison tests, as well as for evaluating the predictive abilities of the linear regression models for cell area and cell speed.

### Metrics used for PCA versus VAE-GAN comparison

Reconstruction accuracy of PCA was evaluated by reconstructing cell contours of the test set using the first ***n*** principal components (where ***n*** = 2, 3, 4, 8, or 16) and programmatically converting them into their corresponding binary masks. Binary cross entropy (BCE) was then used to calculate the mean reconstruction error between all reconstructed and segmented binary masks within the test set. Reconstruction accuracy of the VAE-GAN for latent space dimensions of N = 2, 3, 4, 8, and 16 was evaluated by calculating the mean reconstruction error between the images reconstructed by the decoder and the segmented binary masks, also using BCE, within the test set.

Group separation between PMNs and HL60s in the PCA and latent spaces was evaluated by calculating: 1) the Mahalanobis distances between all pairs of data points within one cell type, and 2) the Mahalanobis distances for all pairs of data points between the two cell types. The non-parametric Wilcoxon rank-sum test was then applied to evaluate the significance of separation between the two groups of distances.

Reproducibility of latent spaces from different training sessions was evaluated using two independent methodologies: 1) Procrustes analysis and 2) correlation analysis of the pairwise distances between data points in any two latent spaces. Procrustes determines the dissimilarity between two sets of points by performing linear transformation (translation, reflection, orthogonal rotation, and scaling) on one set of points to best conform to the same points in the other set by minimizing the sum of their Euclidean distances (36). The correlation analysis involves calculating the pairwise Euclidean distances between all possible pairs of points within one latent space and calculating the correlation coefficient with the pairwise distances of another latent space.

### Calculating cell area and cell speed

Cell area for each segmented image was calculated by summing the total number of pixels that has a non-zero intensity value and normalizing the cell with the largest area to 1. For each time-lapse movie in our data set, we first calculated the migration speed of the cell between two successive time points (a time segment) by dividing the displacement of the cell’s centroid between the two time points with the imaging time interval. We then approximated the instantaneous speed of the cell at each time point by averaging the cell’s calculated speed from the time segments before and after that time point.

### Generating predictive models for cell area and cell speed

Data points with 2, 3, 4, 8, and 16 dimensions in the PCA and VAE-GAN latent spaces were used in predictor variables to fit multiple lasso regression models for the prediction of cell area using 10-fold cross validation, minimizing for root mean squared error (RMS) between predicted and actual cell areas. Comparison of the predictive ability between different models were evaluated using RMS and correlation coefficients between the predicted and actual cell areas of the test set.

Predictive models for cell speed were generated in a similar fashion for VAE-GAN latent spaces (with N = 2, 3, 4, 8, and 16) trained with cell speed encoded in the green channel of the input RGB images. The speed was represented as a centered square with uniform pixel intensity. The intensity for pixels outside of the speed square was set to zero, with the intensity inside the square reflecting normalized cell speed. The number of pixels in the speed square was constant for each image, and was set to be comparable to the number of active pixels across all the binary masks in the data set. This was calculated by performing a pixel-wise summation of all binary masks and grouping pixels with the summed intensities into 50 bins. The total number of pixels, ***p***, with intensities that fall within the 5th and 95th percentiles was then used to construct speed squares of size 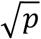 by 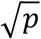 in the speed channel.

---

**Figure S1.**
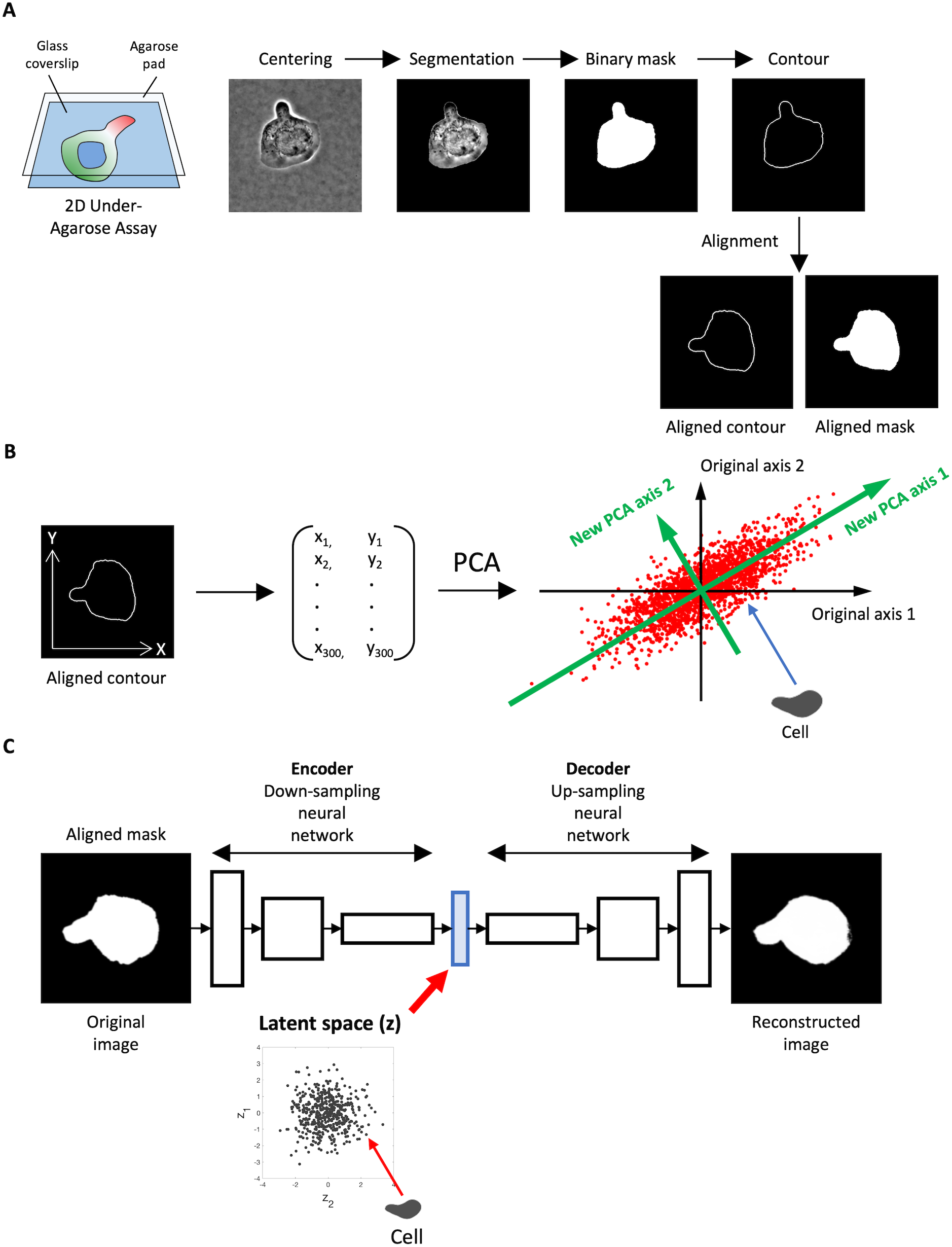
Pipeline for image processing and analysis using PCA and VAE-GAN. (A) Phase contrast images of neutrophils migrating under agarose were first centered and segmented, then converted to binary masks and cell contours, and finally aligned in the +x direction. (B) Aligned contours were parameterized into 600-element vectors (representing 300 points, each specified in x and y) prior to PCA. (C) Aligned binary cell masks were used as training data for a VAE-GAN, to evaluate reconstruction error and mapping into latent space.

**Figure S2.**
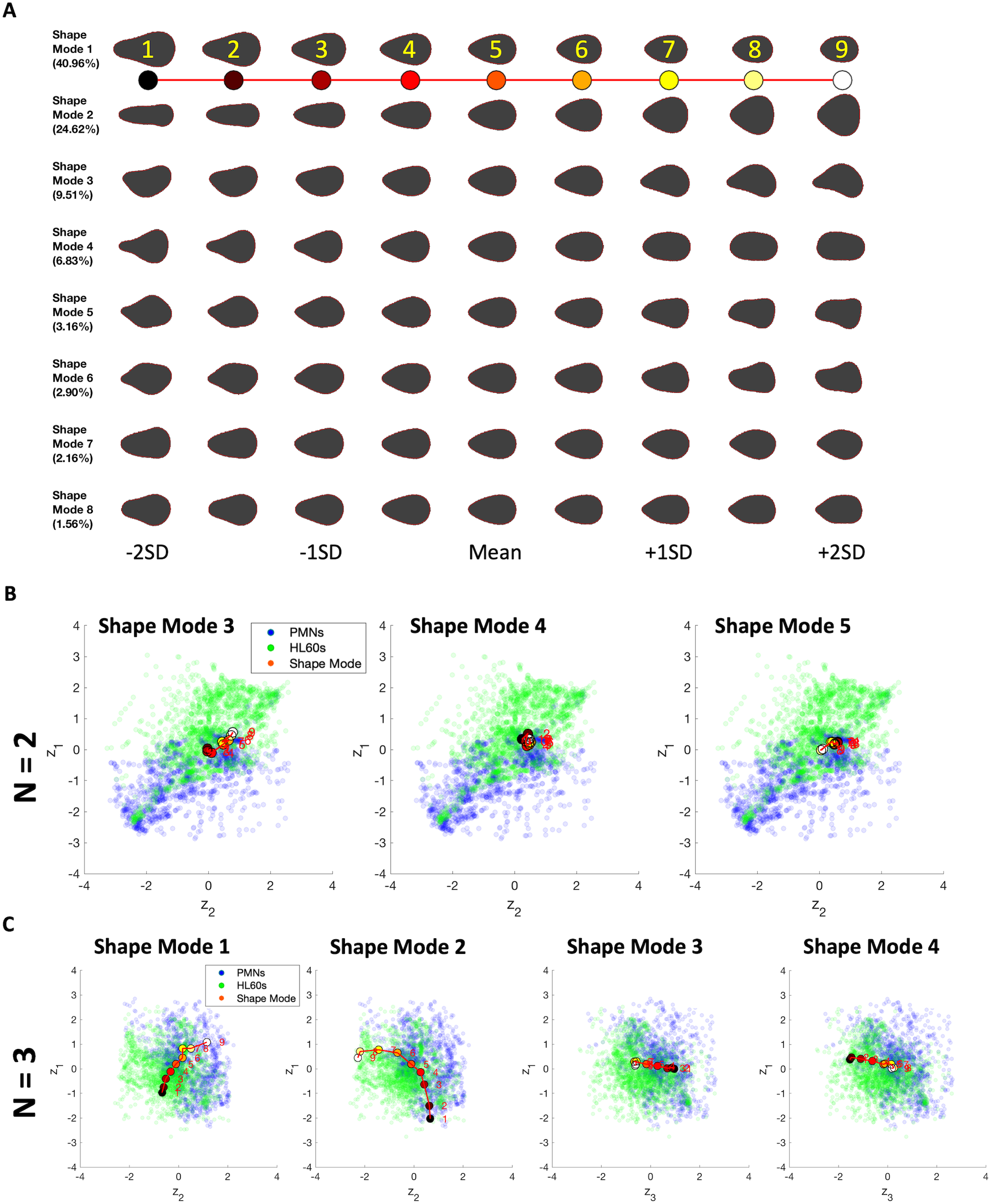
Mapping PCA shape modes to latent spaces. (A) Top 8 principal shape modes determined by PCA showing the numerically-labeled and color-coded reconstructed cell shapes along each shape mode axis. Shape 5 represents the average cell shape for the whole population and is identical for all modes. Shapes 1-4 and 6-9 represent evenly spaced samples along each specified shape mode axis, separated by 0.5 SD. (B) Mapping of principal shape modes 3 to 5 to latent space of N = 2. (C) Mapping of principal shape modes 1 to 4 for latent space of N = 3.

**Figure S3.**
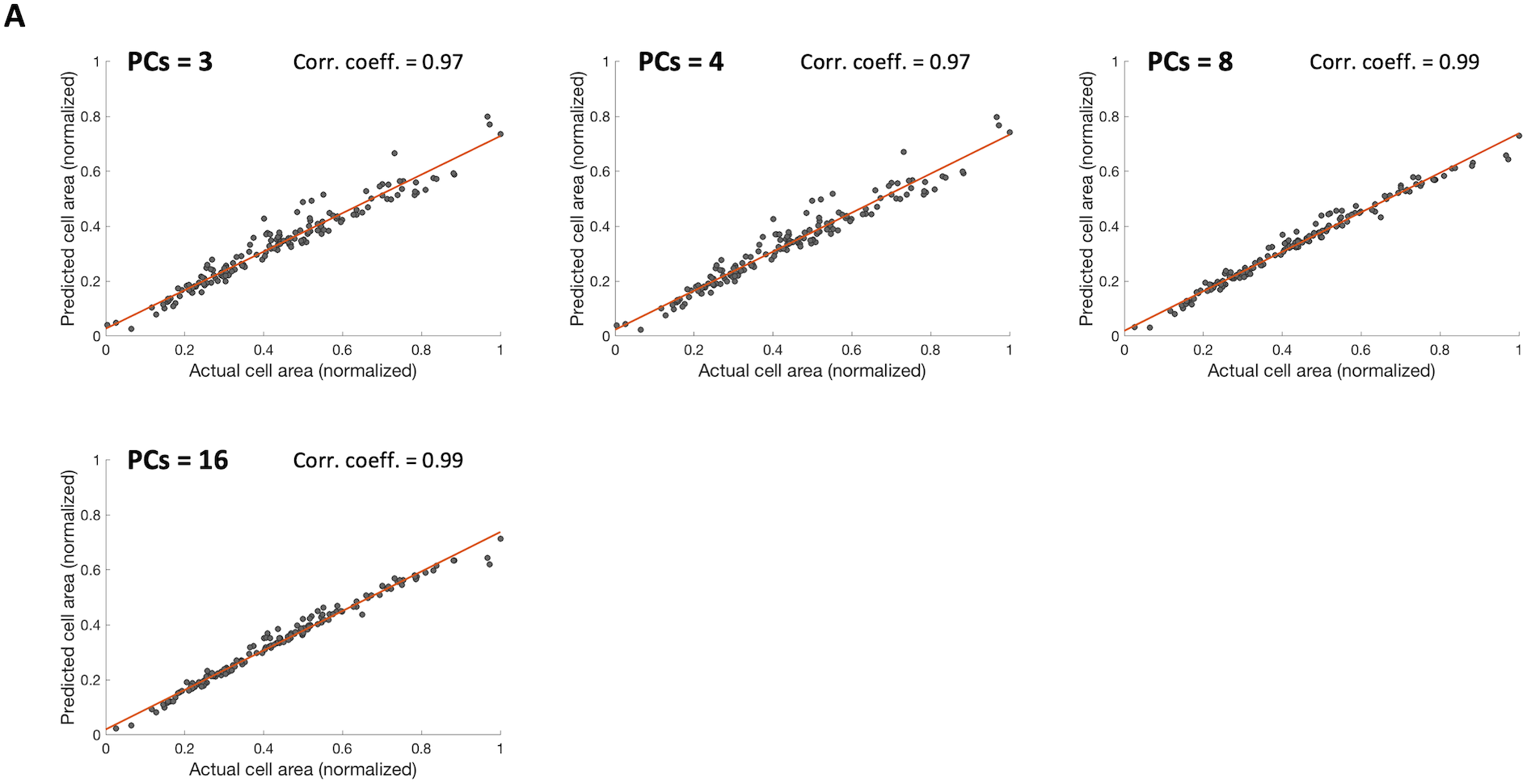
Using PCA shape modes to predict cell area. Predicted versus actual cell area using the first 3, 4, 8, and 16 principal shape modes in separate linear regression models

**Figure S4.**
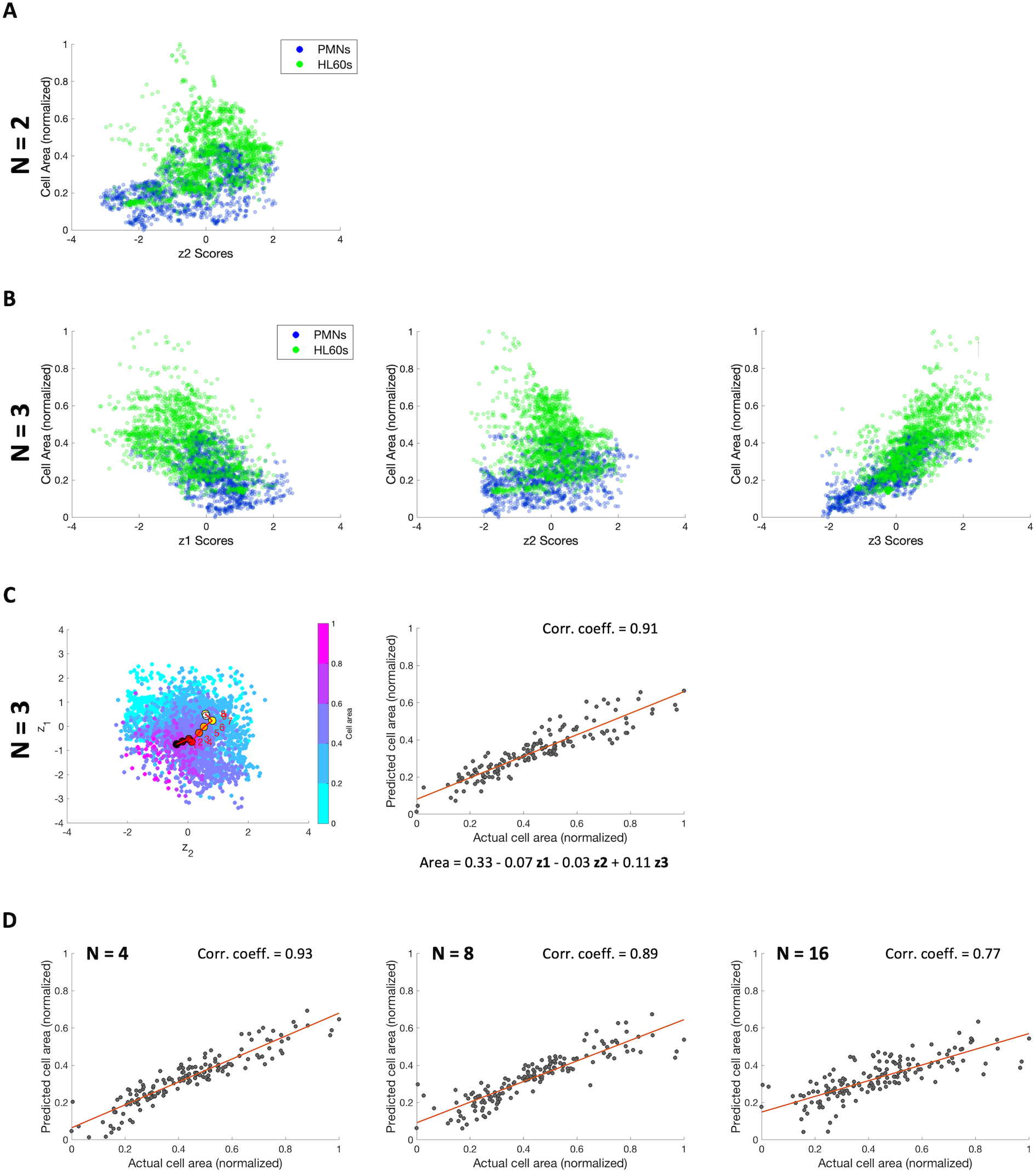
Using latent space embeddings to predict cell area. (A) Scatter plot of cell area versus z_2_ embeddings for N = 2. Blue = PMNs, green = HL60s. (B) Scatter plots of cell area versus embeddings for z_1_ – z_3_ for N = 3. (C) Left, scatter plot of latent space embeddings of z_1_ versus z_2_ for N = 3, color-coded with cell area. Right, predicted versus actual cell area using z_1_ – z_3_ for N = 3 in a linear regression model. Equation shows regression intercept and coefficients for z_1_ – z_3_. (D) Predicted versus actual cell area using all latent space dimensions for N = 4 to 16 in separate linear regression models.

**Figure S5.**
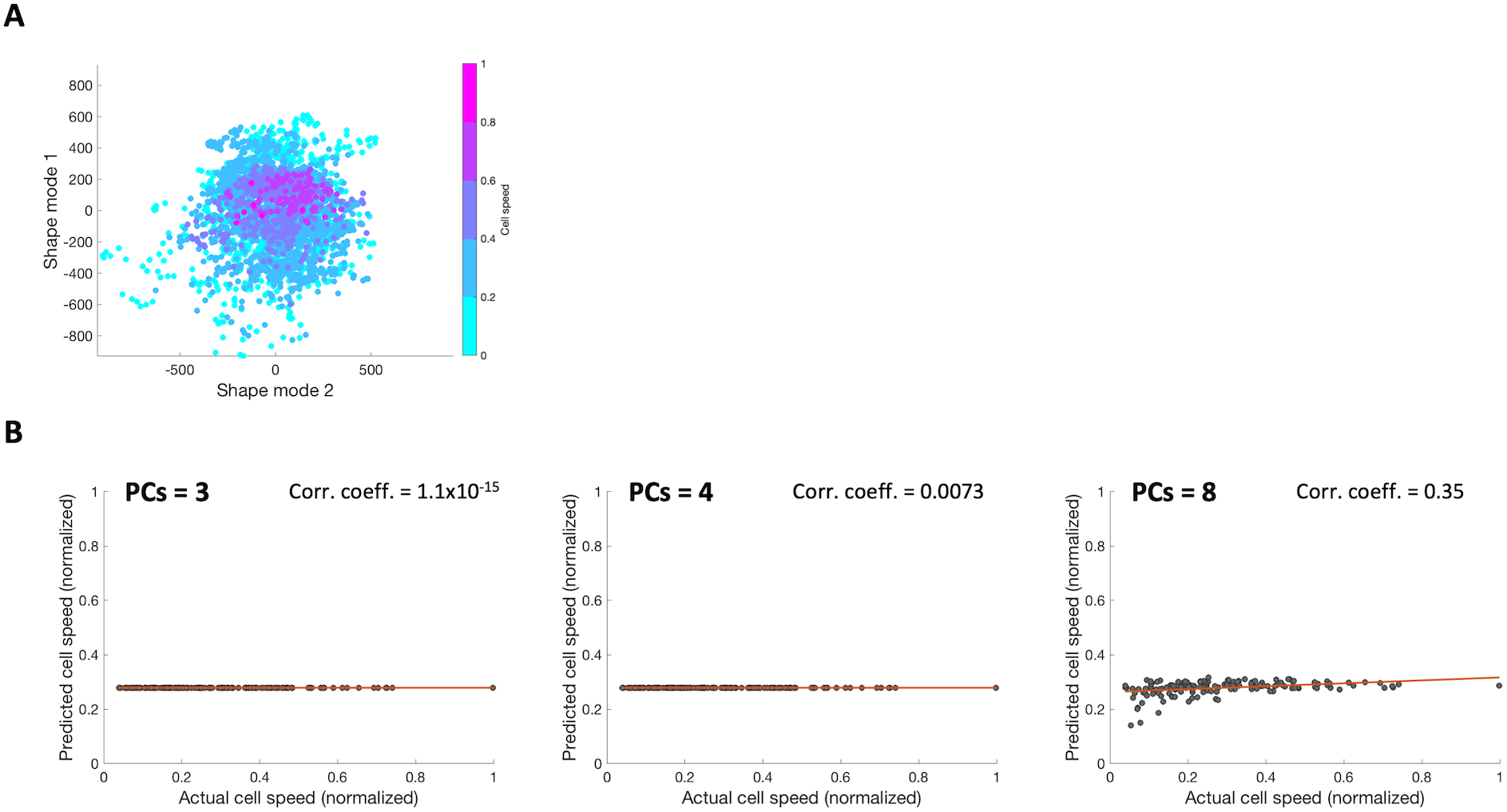
Using PCA shape modes to predict cell speed. (A) Scatter plot showing PC scores for principal shape mode 1 vs. shape mode 2, color-coded with cell speed (B) Predicted versus actual cell speed using the first 3, 4 or 8 principal shape modes in separate linear regression models.

**Figure S6.**
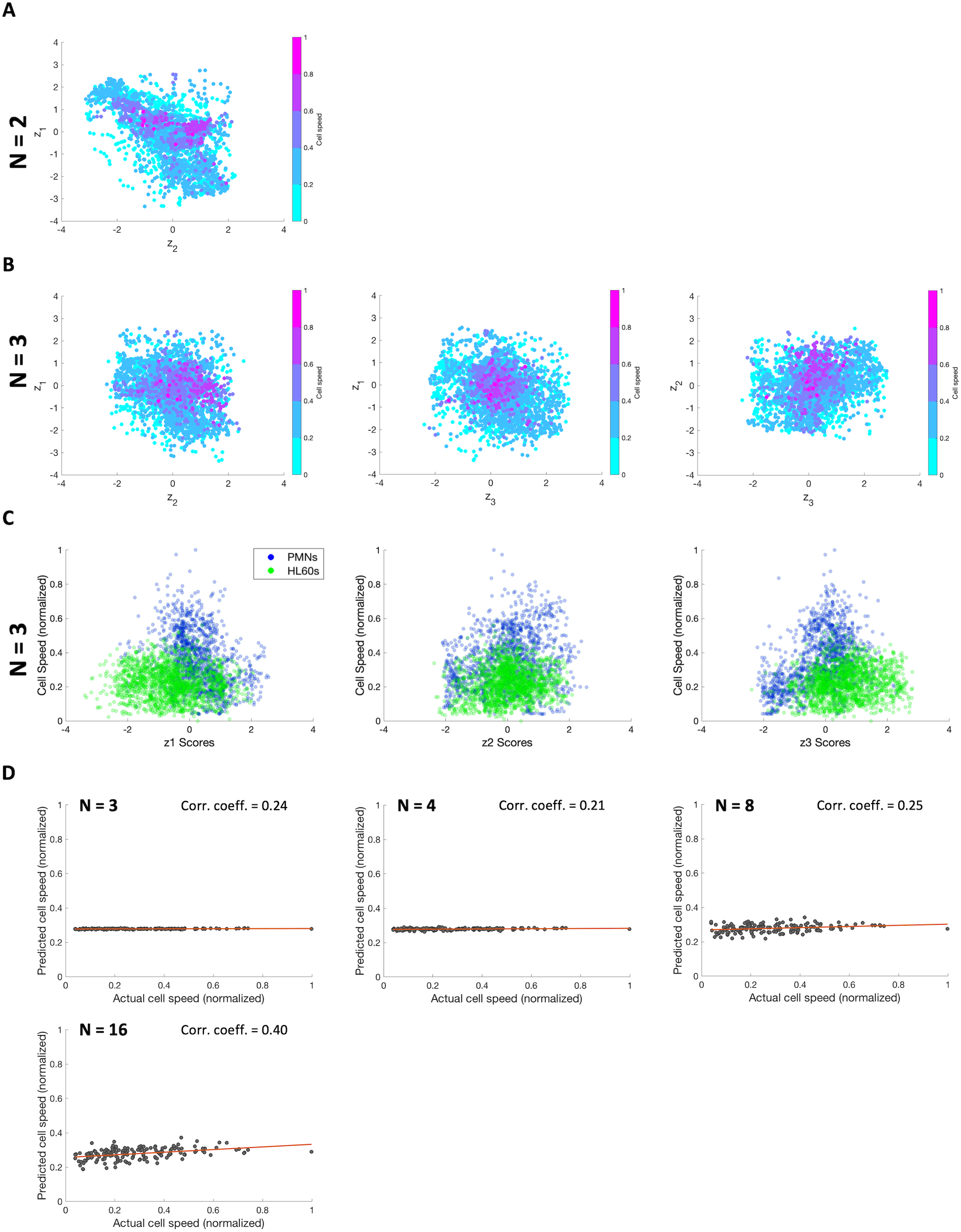
Using latent space embeddings to predict cell speed. (A & B) Scatter plots showing latent space embeddings of z_1_ versus z_2_ for N = 2 and all pairwise dimensions for N = 3, color-coded with cell speed. (C) Scatter plots of cell speed versus embeddings for z_1_ – z_3_ for N = 3. (D) Predicted versus actual cell speed using all latent space dimensions for N = 3, 4, 8 and 16 in separate linear regression models.

**Figure S7.**
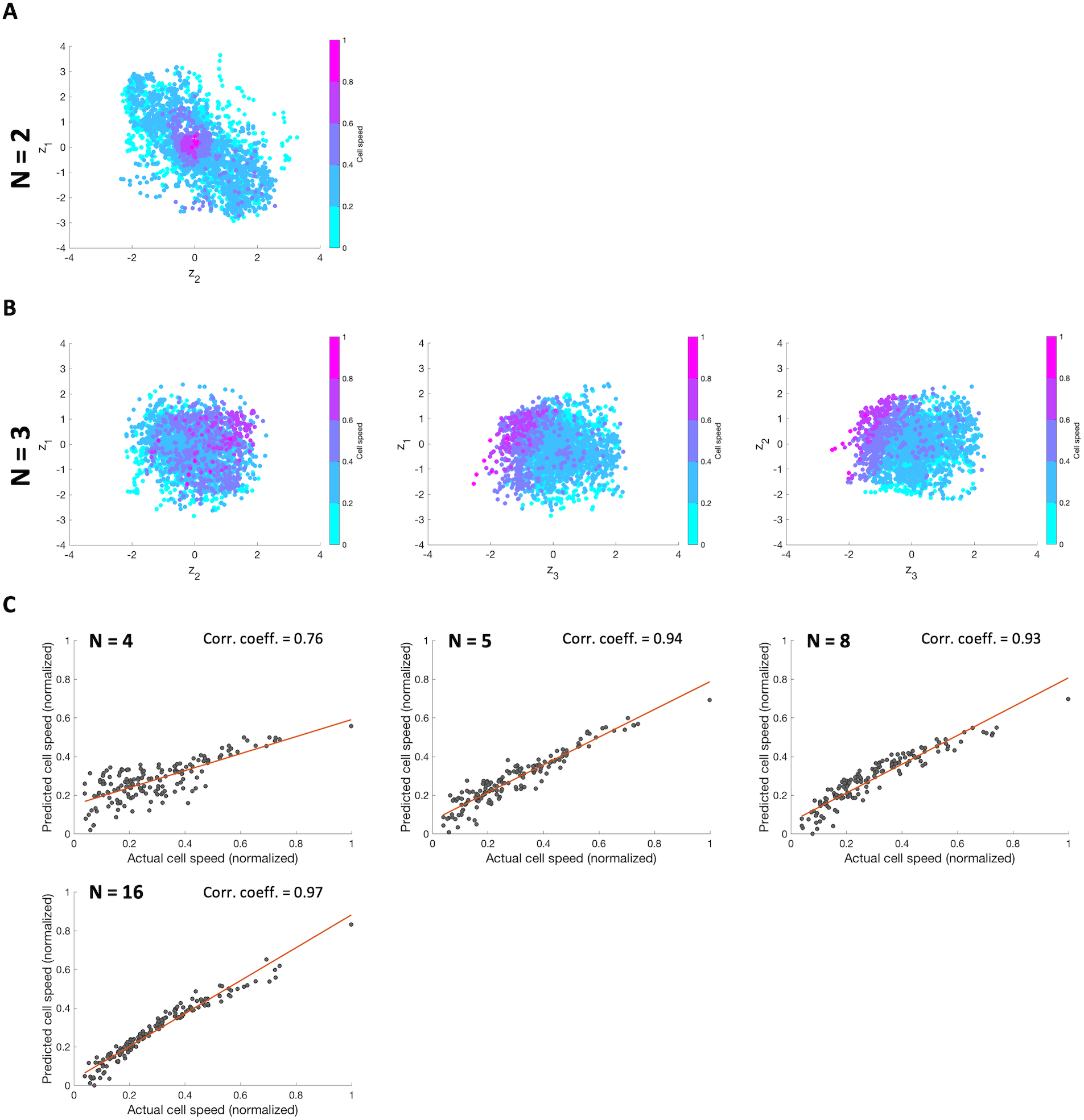
Incorporating cell speed in cell shape analysis using a VAE-GAN. (A) Scatter plot of latent space embeddings in z_1_ versus z_2_ for N = 2, color-coded by cell speed. (B) Scatter plots of latent space embeddings in all pairwise dimensions for N = 3, color-coded by cell speed. (C) Predicted versus actual cell speed using all latent space dimensions for N = 4, 5, 8 and 16 in separate linear regression models.

**Supplementary Movie S1. Time-lapse imaging of PMNs and HL60s**

Representative examples of phase-contrast time-lapse movies for a primary human PMN (left) and a differentiated HL60 cell (right). Both movies have been centered on the centroid of the nucleus (cell frame of reference) and reoriented so that the instantaneous direction of motion is always toward the right. Note the different time intervals for the two movies.

